# Linear Mixed Model of Virus Disinfection by Chlorine to Harmonize Data Collected Across Broad Environmental Conditions

**DOI:** 10.1101/2023.09.12.557160

**Authors:** Mira Chaplin, James Henderson, Kaming Leung, Aleksandra Szczuka, Brianna Hansen, Nicole C. Rockey, Krista Wigginton

## Abstract

Despite the critical importance of virus disinfection by chlorine, we lack a fundamental understanding of the relative susceptibility of different viruses to chlorine and robust quantitative relationships between virus disinfection rate constants and environmental parameters. We conducted a rapid systematic review of virus inactivation by free chlorine and used the resulting data set to develop a linear mixed model that estimates chlorine inactivation rate constants for viruses based on experimental conditions. Over 550 data points were collected in our systematic review, representing 82 viruses over a broad range of environmental conditions. The harmonized inactivation rate constants under reference conditions (pH = 7.53, T = 20 °C, [Cl^-^] < 50 mM) spanned 4 orders of magnitude, ranging from 0.0196 to 1150 L mg^-1^ min^-1^ and uncovered important trends between viruses. Whereas common surrogate bacteriophage MS2 does not serve as a conservative chlorine disinfection surrogate for many human viruses, CVB5 was one of the most resistant viruses in the dataset. The model quantifies the role of pH, temperature, and chloride levels across viruses, and an online tool allows users to predict rate constants for viruses and conditions of interest. Results from the model highlighted potential shortcomings in current USEPA disinfection guidelines.

**Synopsis:** Viruses must be adequately removed by disinfection processes to protect public health. We review data on virus inactivation of chlorine to determine what concentrations of chlorine are required to remove viruses and whether current regulations are adequately protective.

## Introduction

Human and animal viruses play critical roles in human health and well-being. Human noroviruses, for example, cause an estimated 19-21 million annual cases of acute gastroenteritis in the U.S., resulting in tens of thousands of hospitalizations and hundreds of deaths annually^1^ and an annual economic burden of 10.6 billion dollars^2^. Mitigating the spread of mammalian viruses is critical to human health and economics, and the disinfection of air^3^, surfaces^4^, water^5^, and food^6^ is a major route to controlling virus transmission. Free chlorine is the most widely used disinfectant in the United States and is critical for controlling viruses in drinking water and wastewater treatment, as well as in medical^7^ and agricultural applications^8^. Designing effective chlorine disinfection approaches while minimizing disinfection byproducts requires accurate disinfection kinetics data for the viruses and conditions of interest.

The kinetics of virus disinfection by chlorine are highly dependent on water quality parameters. Water pH has a large impact on virus disinfection kinetics because the pK_a_ of HOCl lies close to the typical pH for water disinfection. Hypochlorous acid (HOCl) is a more effective disinfectant than hypochlorite (OCl-); as a result, small changes in pH can substantially alter disinfection kinetics^9^. Temperature also affects virus inactivation rates, with rate constants increasing with increasing temperature^9,10^. The importance of temperature and pH on virus disinfection are reflected in water disinfection guidelines, such as the US EPA’s concentration-time (CT) framework, which require higher CT values for lower temperatures and higher pH values. Other water quality attributes, including chloride concentrations^11–13^ and turbidity^14,15^ also impact virus inactivation rates with chlorine. The very broad range of water types and environmental conditions used in published virus inactivation studies complicates efforts to identify the relative impact of each water quality parameter on virus disinfection and the ways those parameters may interact. The need for harmonized data on virus inactivation with chlorine to update regulations has been noted by the EPA in their regular review of microbial guidelines.^16^

Under identical environmental conditions, inactivation rates differ greatly between viruses.^13,17–23^ For example, CT values for 3-log_10_ virus inactivation are over 1000x greater for coxsackievirus B5 (CVB5) than for murine norovirus or human adenovirus 2^23^. Order-of magnitude differences in inactivation rates also exist even for viruses of the same species.^24,25^ CT values for 2-log_10_ inactivation of distinct viruses belonging to the species *Enterovirus B* varied from 0.82 to 3.6 mg*min/L,^26^ and some CVB5 environmental isolates have rate constants up to five times lower than the laboratory strain CVB5 Faulkner.^24^ Mechanistic studies on closely related viruses have identified specific virus characteristics, such as genome and protein composition, that contributed to these differences in kinetics^27,28^ but it is not known whether those characteristics have universal impacts across viruses. A better understanding of the relative susceptibility of different groups of viruses to chlorine would help identify more general virus inactivation mechanisms and help in the selection of surrogate viruses for chlorine disinfection processes.

Models are frequently used to fill in gaps in the experimental disinfection data, as it is not practically feasible to test the susceptibility of every virus to free chlorine under every relevant condition. For example, data collected within individual studies have been incorporated into models that predict chlorine inactivation for human adenovirus 2 and CVB5 with water quality parameters as inputs.^9,29^ More recently, Kadoya et al. used reviewed literature data to build predictive models for virus disinfection as a function of pH, temperature, assay, and water type.^30^ Their models focused on select groupings of human viruses and identified differences within virus species (e.g., different strains of poliovirus), rather than between virus species or families. As the virus disinfection literature encompasses a broad range of viruses and environmental conditions, an opportunity exists to build robust statistical models to identify important trends between broad virus groupings and predict inactivation kinetics under a range of environmental conditions.

In this study, we conduct a rapid systematic literature review to generate a dataset of high-quality chlorine disinfection rate constants for a broad range of human and non-human viruses. We then apply these data to build a linear mixed mode to identify major trends in susceptibility between groups of viruses (e.g., poliovirus versus MS2 bacteriophage) and quantify the roles of environmental conditions and laboratory-specific impacts (i.e., random effects) on virus disinfection rates. Furthermore, the model enables prediction of the inactivation rates of studied viruses under specific environmental conditions that are not reported in the literature. The resulting data and models are valuable resources for the widespread application of chlorine disinfection, will facilitate the selection of conservative surrogates for viruses of concern, and will identify research gaps that can move us closer to predicting chlorine inactivation of any virus, including emerging and nonculturable viruses, under conditions relevant to water treatment.

## Methods

### 2.1 Systematic Review

We conducted a systematic review on viral inactivation by free chlorine, according to the guidelines published by the Preferred Reporting Items for Systematic Reviews and Meta-Analyses (PRISMA)^31^. We set criteria to ensure that chlorine concentration and virus infectivity were accurately reported and chlorine concentrations were relevant for water and wastewater treatment systems. Articles were included for data extraction when they fit the following criteria: virus removal was assessed using culture-based assays; temperature and pH were reported; chlorine decay was described or the study was conducted with experimental protocols sufficient to minimize chlorine decay; the study was conducted on non-aggregated viruses in solution; the study was conducted in pure water, buffers, natural waters or wastewater that had undergone secondary treatment; data reported included at least one time point under five minutes; the chlorine concentration was less than 20 mg/L; sufficient kinetic data was reported to extract second-order rate constants; the study was performed the study on a virus with a publicly available genome. Additional details of the review are provided in Supplementary Information S1.

### 2.2 Rate Constant Extraction

Two reviewers extracted all raw data using Digitizeit software^32^ and independently calculated second-order rate constants. A number of studies reported second-order rate constants; however, to facilitate direct comparison, we calculated the rate constants from the raw virus inactivation data whenever it was available. In cases where inactivation exhibited tailing after 2- to 4-log_10_ inactivation, reviewers only extracted data from the initial inactivation phase. Second-order rate constants were calculated for each dataset based on the Chick-Watson model:

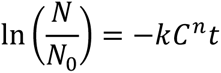

where t is exposure time of the virus to chlorine, N is the concentration of infectious virus at time t, N_0_ is the concentration of infectious virus at time zero, k is the second-order rate constant, and C^n^ is chlorine concentration to the power of the coefficient of dilution. We assumed n was equal to 1 to enable direct rate constant comparisons. The units of C^n^ t are therefore min*(mg/L), and the term is equivalent to the CT value. This results in the following equation to calculate rate constants:

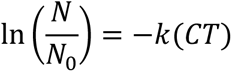

This equation was used to determine the second-order rate constant k with units of L mg^-1^ min^-1^, as described in Supplementary Information S2.

We assessed agreement between reviewers for each data point using the Lin’s concordance coefficient. Inactivation rates presented and used for modeling are the mean of the values calculated by two reviewers.

### 2.3 Model Variable Development

We extracted details on experimental conditions, virus characteristics, and study authors from the manuscripts and online databases and ultimately used several of these as variables in our models (Table S2). To assess agreement between reviewers, experimental condition variables were checked by a second reviewer for all data points from 20 of the 72 papers. All examined experimental variables were identical between reviewers. The variables considered in the model either have been associated with chlorine inactivation in the literature or were identified via residuals analysis. Brief descriptions of the variables are provided below, and additional assumptions and justification are outlined in the Supporting Information S3.

#### Temperature

Temperatures extracted from the literature were converted to degrees Celsius. We set temperatures to the mean of provided ranges and assumed that room temperature was 20 °C when not defined. For modeling, temperature was referenced to 20 °C in increments of 5 °C.

#### pH

We extracted pH values from manuscripts and referenced pH values to 7.53, the pK_a_ of HOCl. To calculate ΔpH^2^, we squared the difference between the reported pH value and 7.53. We calculated α_HOCl_, the proportion of the free chlorine in the form of HOCl, as a nonlinear function of pH using the equation α_HOCl_ = [H+]/([H+] + K_a_) where [H+] is the concentration of H+ ions, and K_a_ is the acid dissociation constant.

#### Water type

We divided water type used in studies into four categories: buffer solutions, low organics natural or treated waters, high organics natural or treated water, and ultrapure water.

#### Chloride content

We included a Boolean variable labeled “high chloride” that represented chloride concentrations that met or exceeded 50 mM (1773 mg/L Cl-). We chose this threshold because most studies fell either between 0 - 10 mM Cl- or >100 mM Cl-. When solution composition was not available, we assumed that solutions were low chloride.

#### Virus purity

As virus purification minimizes chlorine decay in inactivation experiments^33^, we defined purification level as an experimental variable. Purification level was divided into three categories: high, medium, and low. High was defined as gradient purification or equivalent, low purification was defined as no purification or only low levels of centrifugation, and medium purification included purification steps between those two levels (e.g., chloroform extraction).

#### Virus classification

The Baltimore classification system divides viruses based on the steps of mRNA production, and incorporates information about the genetic material (DNA or RNA), whether a virus is single or double stranded, the sense of a single-stranded RNA virus, and whether a reverse transcription step occurs in virus reproduction.^34^ As data were not found for Baltimore classes using reverse transcription (VI and VII), Baltimore classes are referred to by their sense, strandedness, and genetic material (e.g. (+)ssRNA for positive-sense single-stranded RNA). An additional Boolean variable, “DNA” was created, DNA viruses labeled as 1 and RNA viruses labeled as 0. Taxonomic information on family, genus, and species was extracted from the International Committee on Taxonomy of Viruses (ICTV) and is based on the 2021 taxonomic release.^35^

#### Virus name

ICTV does not classify viruses beyond the species level, although subspecies have unique biological and clinical characteristics (e.g., human norovirus and murine norovirus are in the same species). We therefore included a variable for “virus name” that was defined as the most specific taxonomic unit available.

#### Virus Morphology

Virus features were identified for the viruses included in the extracted data set. These features include particle diameter, the presence of an envelope, the presence of a tail, genome length, and virus symmetry (helical vs icosahedral).

#### Other study data

Each article was reviewed to determine the corresponding author and year of publication. When no corresponding author was designated, we assumed a corresponding author based on author descriptions and the location where experiments were performed.

### 2.4 Model Development

We generated a linear mixed model to probe the relative impact of experimental conditions, virus taxonomy and physical properties, and within-paper variation (e.g., systematic effects on rate constants attributable to being from a certain paper) on virus inactivation by free chlorine. The Akaike Information Criterion (AIC) was used to assess model fit. The AIC penalizes each parameter, so only independent variables that improved the log likelihood enough to overcome the imposed penalty were accepted into the model. Conditional R^2^ values were also included for comparison purposes. R^2^ values were calculated using MuMin package^36^ which bases R^2^ values on equations developed by Nakagawa et al.^37,38^

We fit models to the following equation:

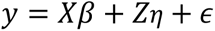

where y is a n-vector of the log_10_ inactivation rates, X is an (n × p) feature matrix of independent variables with n equal to the number of samples and p to the number of features, *β* is a vector of model coefficients, Z is a random effect design matrix with columns identifying inactivation rates from the same paper (or, in some models, the same corresponding author), *η* is a vector of Gaussian random effects with mean 0, and *ϵ* is a vector of model residuals.

Beginning with a model including only virus species, we used a stage-wise forward model selection procedure to determine which variables to include based on AIC (Supporting Information S4). To start, we ran a model with virus name as the only feature (Figure S3, M0). We next generated a minimal model, identified as M1, that included virus name, temperature, and pH as independent variables. We accounted for within-paper correlations using a per-paper random intercept and compared this to a model using random intercepts for corresponding authors. Specifically, we included publication ID and corresponding author both independently and jointly as random effects in this model and each updated model with additional features. We then considered for inclusion additional features including buffer, purification level, α_HOCl_, year of publication, and high chloride. Finally, given previous research showing that the interaction between chloride and pH could be significant to virus inactivation,^39^ we tested the interaction of high chloride and pH terms. We conducted residuals analysis and tested additional features and interactions suggested by patterns in the residuals, including ΔpH^2^ and the interaction between temperature and Baltimore class (Figure S4).

Due to variable collinearity, a linear mixed model of this dataset can include only one virus feature, namely a taxonomy-related or structure-related independent variable. To test the effect of different virus taxonomic units and virus characteristics on the AIC without introducing confounding variables, we replaced virus name in our optimized model with independent variables for species, genus, family, Baltimore class, diameter, genome length, tail, or symmetry (Figure S5). The analyses described above were performed in R 4.2.1 using the lme4 package.^40^ The data files and the scripts for data processing, model development, and figures are available in GitHub at https://github.com/mira-create/chlorine_model.

## Results and Discussion

### 3.1 Rapid Systematic Review

We identified 1,366 unique articles in the rapid systematic review of rate constants for virus inactivation by free chlorine. Of these, 497 underwent full-text review and 72 of those were included in the final dataset, resulting in 564 extracted chlorine disinfection rate constants. Articles were most commonly excluded due to insufficient characterization of initial chlorine concentrations or chlorine decay, chlorine concentrations that were far higher than the 1-20 mg Cl_2_/L typically used during drinking water and wastewater treatment,^41^ or a lack of infectivity measurements (Figure S1). The rate constants extracted by the two reviewers had a Lin’s concordance coefficient of 0.997 (95% CI: .995-.999) (Figure S2). The two independent reviewers calculated rate constant values within ten percent of each other for 534 of the 564 rate constants. The >10% difference in the remaining 30 rate constants was due to a lack of plot resolution in the reviewed studies.

The resulting inactivation data represent 82 viruses from 18 families and five of the seven recognized Baltimore classes (Figure 1). The data contain no rate constants from Baltimore classes VI (retroviruses) or VII (hepadnaviruses). In total, 72% (406) of the extracted rate constants were for (+)ssRNA viruses, followed by dsDNA viruses (88, 16%), dsRNA viruses (43, 8%), ssDNA viruses (20, 3.5%), and (-)ssRNA viruses (7, 1.2%). The most studied groupings are common human viruses that can be transmitted via water and common bacteriophage surrogates for human viruses. The specific viruses with the greatest number of reported rate constants were poliovirus 1 Mahoney (86), MS2 bacteriophage (46), and Human Adenovirus 2 (46).

**Figure 1.**
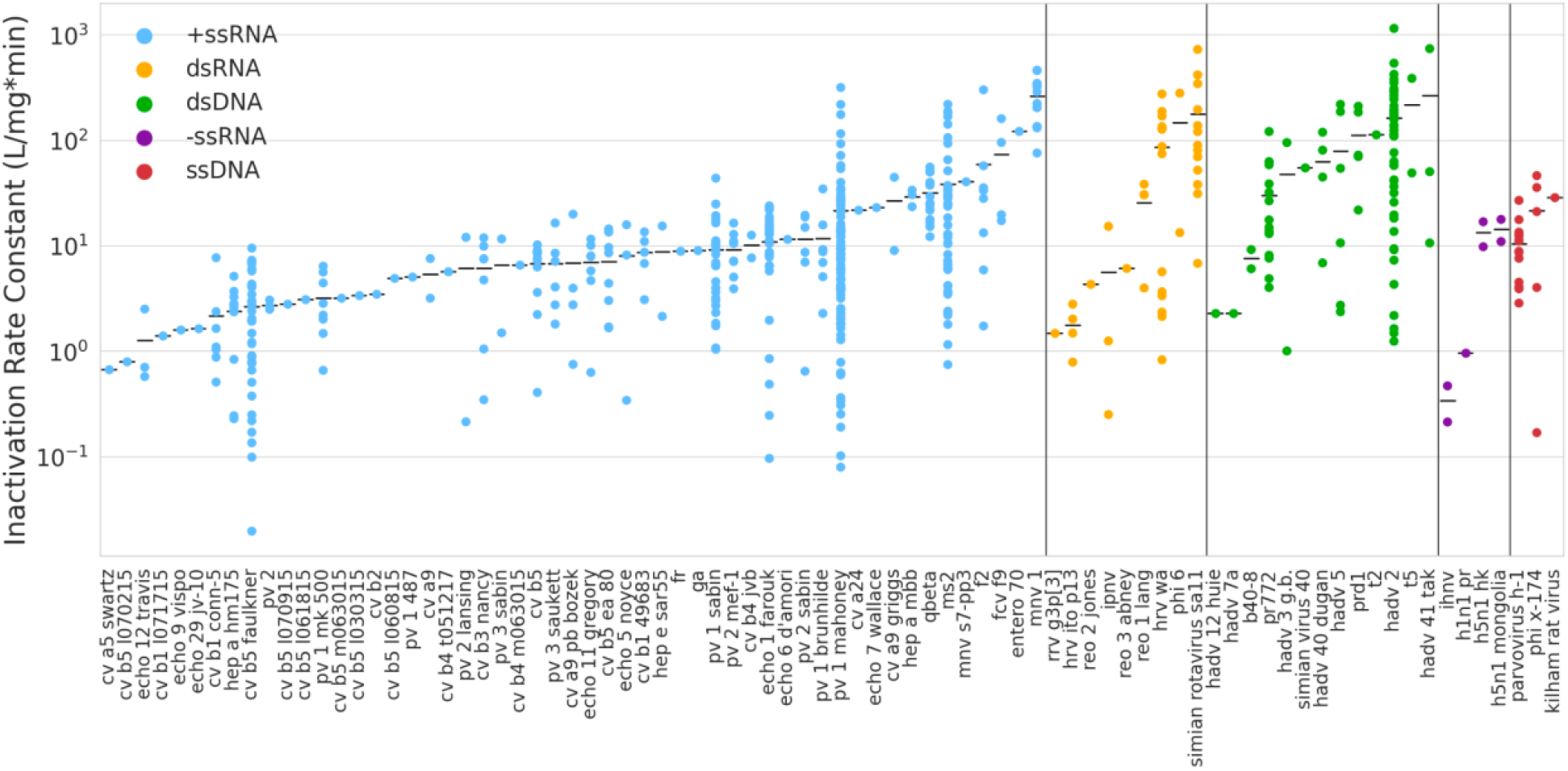
Inactivation rate constants under all experimental conditions calculated from the systematic review, grouped by Baltimore class. Within each Baltimore class, viruses are sorted by average inactivation rate. The y axis represents the magnitude of the inactivation rate constants on a log_10_ scale.

The dataset is missing notable groups of viruses and data at specific water conditions that are relevant for disinfection processes. For example, the dataset lacks several of the viruses that the World Health Organization (WHO) has identified as of concern for waterborne transmission,^42^ including astroviruses, sapoviruses, parechoviruses, and hepatitis E virus (Table S4). The widespread use of chlorine as an all-purpose disinfectant underlines the need for high-quality chlorine rate constants for viruses that are spread beyond the fecal-oral route. In this regard, there are few rate constants for human respiratory viruses such as influenza viruses (5 rate constants) or coronaviruses (0 rate constants). Likewise, viruses that disrupt agricultural and food processing operations, such as livestock viruses, plant viruses, fish viruses, and dairy phages are also largely absent. Some of these identified gaps in viruses are due to the absence of cell culture assays (e.g., Parechoviruses^43^) and recent culture method advances ^44,45^ could facilitate the collection of high-quality disinfection data for additional viruses. Dividing the Baltimore class data by pH, temperature, and chloride level highlights the uneven data across experimental conditions (Figure S7). Whereas the (+)ssRNA viruses have broad coverage across relevant pH, temperature, and chloride conditions, the other virus classes are poorly represented across certain conditions. For example, there was no or limited data collected under high chloride conditions and for pH values above 8 for a number of Baltimore classes, and environmental temperature ranges are poorly represented for (-)ssRNA.

The calculated second-order inactivation rate constants of all data span six orders of magnitude, from 0.0196 to 1150 L mg^-1^ min^-1^. Rate constants for individual viruses varied greatly across studies. For example, inactivation rate constants for poliovirus 1 Mahoney ranged from 0.0792 to 315 L mg^-1^ min^-1^. High variation in reported inactivation rate constants likely stems from the wide range of experimental conditions encompassed in the data, especially broad temperature and pH ranges. Efforts to narrow down experimental conditions, however, led to drastic decreases in the size of the data set (Figure S8). When the dataset was narrowed to include rate constants collected at approximately room temperature (15 -25 °C) and pH values between 6 and 7, the number of viruses represented in the data decreased from 82 to 28 and the number of rate constants decreased from 564 to 94. Surprisingly, narrowing the conditions did not decrease the rate constant variance for an individual virus; for example, the rate constant standard deviations for MS2 bacteriophage and poliovirus 1 Mahoney increased from 54.5 to 69.1 L mg^-1^ min^-1^ and from 46.4 to 56.4 L mg^-1^ min^-1^, respectively. This indicates that more complex modeling is required to quantify trends between viruses and environmental conditions.

### 3.2 Model Selection

We evaluated successive model improvement using AIC and the full sequence of models is presented in Supporting Information section S4. The stagewise forward model selection resulted in the following equation for a data point i collected in publication p(i) for virus v(i) (AIC = 582, R^2^ = .866):

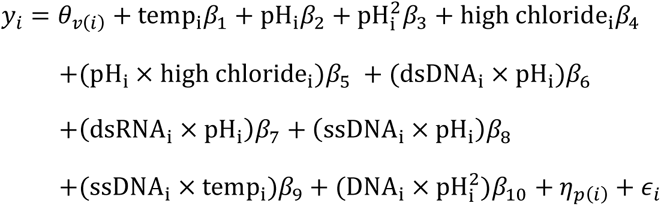

where y_i_ is the log_10_ of the inactivation rate, *β* values are the model coefficients, *η*_p(i)_ is the random effect coefficient for the publication, and *ϵ*_i_ is the model residual. This model includes an intercept, temperature, high chloride, pH, ΔpH^2^, and several interactions as independent variables. The model also includes publication ID as a random effect.

The incorporation of temperature, pH, chloride concentration, and the interaction between pH and chloride all improved the model, confirming their importance in chlorine inactivation of viruses.^9–12,39,46^ Water type and virus purification level did not improve the model despite having previously been shown to impact disinfection by chlorine in individual studies.^33,47,48^ This is likely due in part to the broad classifications of these variables used in the model. For example, for water type, buffer solutions and natural/treated waters encompass waters with a range of characteristics. More granular classification, however, risks overfitting the model due to the small numbers of data points in some categories. The addition of a publication-specific identifier as a random effect improved the model, however the addition of corresponding author as a second random effect did not lead to further improvement. This indicates that data points from different papers written by the same corresponding authors are less correlated than data points from the same paper. Of the two nonlinear functions of pH tested, ΔpH^2^ improved the model, whereas α_HOCl_ (calculated using [H+] concentration) did not. The improvement generated by ΔpH^2^ indicates nonlinear effects on inactivation relative to the pK_a_, while the lack of impact of the α_HOCl_ may be due to the role that both HOCl and OCl-play in virus inactivation.

Five additional interaction terms appear in the final model, namely dsDNA and pH, dsRNA and pH, ssDNA and pH, ssDNA and temperature, and DNA and pH^2^ (see Supporting Information S4 for details). This suggests that the rate constant predictions are impacted differently by temperature and pH. It is possible that the effects of experimental conditions on inactivation constants are impacted by smaller taxonomic units than Baltimore class, such as virus family, genus, species, etc. However, the current dataset is insufficient to test if the model improves by including interactions between pH or temperature and smaller taxonomic units.

As discussed above, only one virus attribute can be included in any model due to variable collinearity. Once the experimental features were selected, we tested the effect of different virus taxonomic units and structural characteristics in lieu of virus name on the AIC (Figure S5). All tested independent taxonomic and structural variables (i.e., tail, capsid structure, diameter, genome length, Baltimore class, family, genus, and species) lowered the AIC and improved the model except for lipid envelope. Features related to virus taxonomy had greater AIC effects than structural features, likely due to the highly uneven dataset for structural features. For example, only 9 of the 564 data points corresponded to enveloped viruses. Of the taxonomic and structural features, we selected virus name in the model because it produced the lowest AIC.

### 3.3 Model Results

#### Model Predictions Under Reference Conditions

Whereas the systematic review highlighted the limited number of viruses with rate constants collected under similar experimental conditions, the model enables harmonization of rate constants for all viruses included in the systematic review. We use the term harmonize to describe the use of models to provide a comparable view of data from different sources collected under different conditions. Under reference conditions (pH = 7.53, T = 20 °C, [Cl^-^] < 50 mM), the predicted inactivation rate constants for the 82 viruses identified in the systematic review covered four orders of magnitude (0.510 L mg^-1^ min^-1^ to 554 L mg^-1^ min^-1^; Supplementary Information section S11), two orders of magnitude smaller than the rate constant range observed from the systematic review results that covered all conditions (0.0196 to 1150 L mg^-1^ min^-1^). The confidence intervals generated by the model are narrower for viruses that had been more widely studied under a wide range of experimental conditions. For example, the confidence intervals for the reference condition rate constants of widely-studied MS2 bacteriophage (95% CI: 7.76-18.3) and poliovirus 1 Mahoney (95% CI: 4.05-10.7) are relatively small. In contrast, the confidence intervals for the reference condition rate constants for influenza A/WhooperSwan/Mongolia/244/2005 (95% CI: 2.91-254) and rhesus rotavirus G3P[3] (95% CI: 0.130-16.5) were relatively large, as each virus was included in a single publication. Large confidence intervals limit the ability to determine if one virus is more resistant to inactivation by chlorine than another; only 36 out of the 81 (44.0%) viruses in the model have predicted inactivation rate constants under reference conditions that differed statistically (with nominal p < .05) from the model reference virus MS2 bacteriophage.

Despite these large confidence intervals, the harmonized virus rate constants do exhibit trends between certain virus groupings. The (+)ssRNA viruses and dsDNA viruses are of particular interest for water treatment due to a number of important waterborne viruses in these classes. The harmonized dataset suggests dsDNA viruses are generally more susceptible to free chlorine than the (+)ssRNA viruses (Figure 2a), which may be related to the larger sizes of the dsDNA viruses (Table S6). Previous research has implicated protein damage in viral inactivation by free chlorine^27^ and larger viruses have more protein targets for reactions with chlorine compared to smaller viruses.

**Figure 2.**
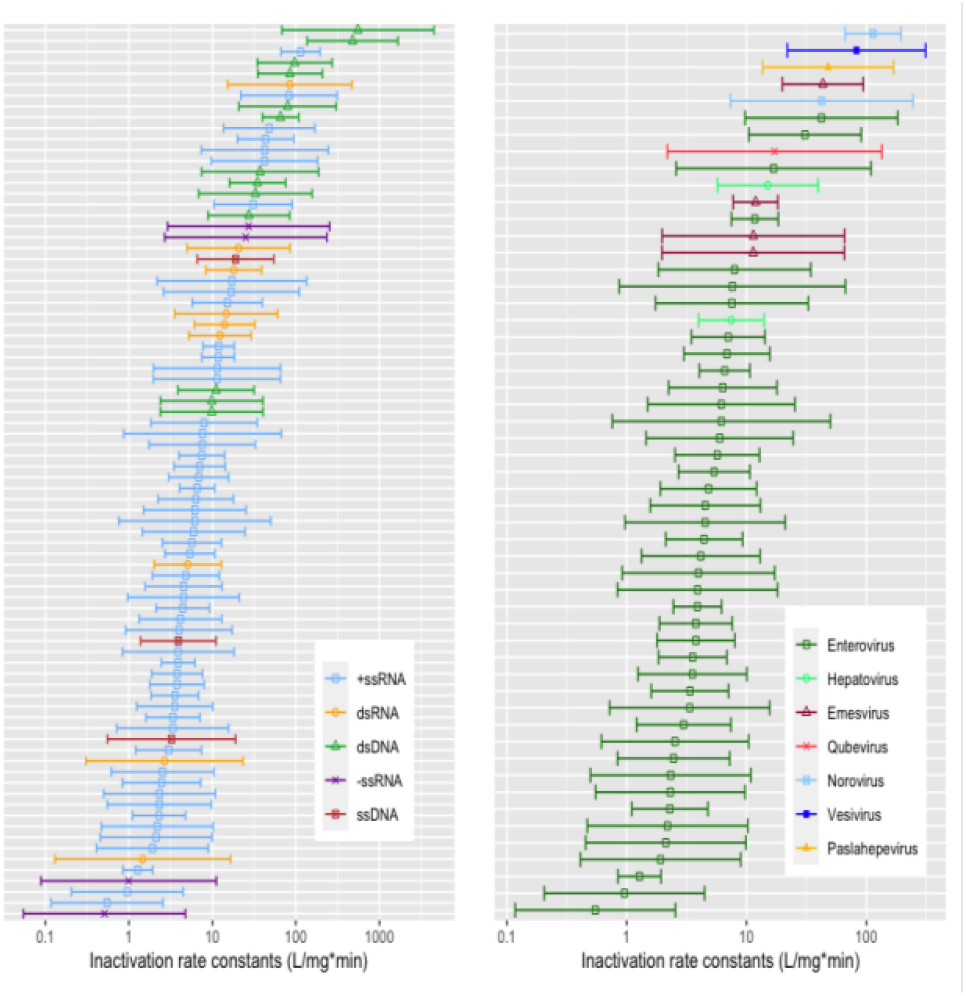
Harmonized rate constants generated by the model under reference conditions (pH = 7.53, T = 20 °C, [Cl-] < 50 mM). a) Chlorine inactivation rate constants for all viruses with experimental data extracted in systematic review, colored by Baltimore class. b) Chlorine inactivation rate constants for all (+)ssRNA viruses with experimental data extracted in systematic review, colored by virus genus. Error bars represent 95% confidence intervals of the rate constant predictions. See Supporting Information section S11 for predicted rate constants at reference conditions.

The (+)ssRNA viruses exhibited a broad range of chlorine susceptibilities (Figure 2b). In particular, the (+)ssRNA enteroviruses, members of the most highly represented virus genus in our dataset, exhibited rate constants that ranged nearly three orders of magnitude at reference conditions, from 0.558 L mg^-1^ min^-1^ (95% CI: 0.117-2.56) for CVB5 l070215 to 42.0 L mg^-1^ min^-1^ (95% CI: 9.75-183) for enterovirus 70. Several enteroviruses had the lowest rate constants overall at reference conditions, including CVB5 Faulkner and poliovirus 1 MK 500. Previous research on enterovirus inactivation by free chlorine linked chlorine disinfection susceptibility to the presence and solvent accessibility of cysteine, methionine, and tyrosine residues, which could explain the wide variation across enteroviruses.^24,28^ While the (+)ssRNA human norovirus was not represented in our dataset, three related viruses that are used as human norovirus surrogates (murine norovirus 1, murine norovirus s7-pp3, and feline calicivirus f9) all had higher predicted inactivation rate constants at reference conditions than all enterovirus rate constants. These (+)ssRNA virus results suggest that human noroviruses will be effectively disinfected when chlorine treatment is designed to disinfect the conservative human enteroviruses. Future research should expand the diversity of (+)ssRNA viruses tested with free chlorine to confirm if the large range of chlorine susceptibilities exhibited by enteroviruses is typical of other virus genera.

Our dataset suggests there are other viruses with inactivation rates at reference conditions in the range of the most resistant enteroviruses, below around 3 L mg^-1^ min^-1^. These include (-)ssRNA infectious hematopoietic necrosis virus (ihnv), dsRNA rhesus rotavirus g3p, ssDNA kilham rat virus, and ssDNA h1 parvovirus. Many of these viruses have not been included in other systematic reviews that focused primarily on common human waterborne viruses^30,49^. Future research should probe these viruses further to identify the specific contributing traits to high chlorine resistance.

#### Model Quantifies Role of Experimental Conditions

Beyond harmonizing the data, the model quantifies overarching trends with pH, temperature, and chloride conditions. Temperature, pH, and chloride trends agree with previous findings with individual viruses, namely that inactivation rate constants increase with increasing temperature, decrease with increasing pH, and increase with increasing chloride levels (Figure 3). For all Baltimore classes except ssDNA, every five degree increase in temperature leads to a predicted increase in inactivation rate constant of 0.0967, or a 24.9% increase (95% CI: 18.8%-31.4%). Due to the multiple model terms involving pH, different classes of viruses behave differently at different pH values. For MS2 bacteriophage, a 1 unit increase in pH from 7.53 to 8.53 under low chloride conditions and at any temperature leads to a predicted decrease in inactivation rate constant of 0.216, or 39.2% (95% CI:30.8%-46.6%).

**Figure 3.**
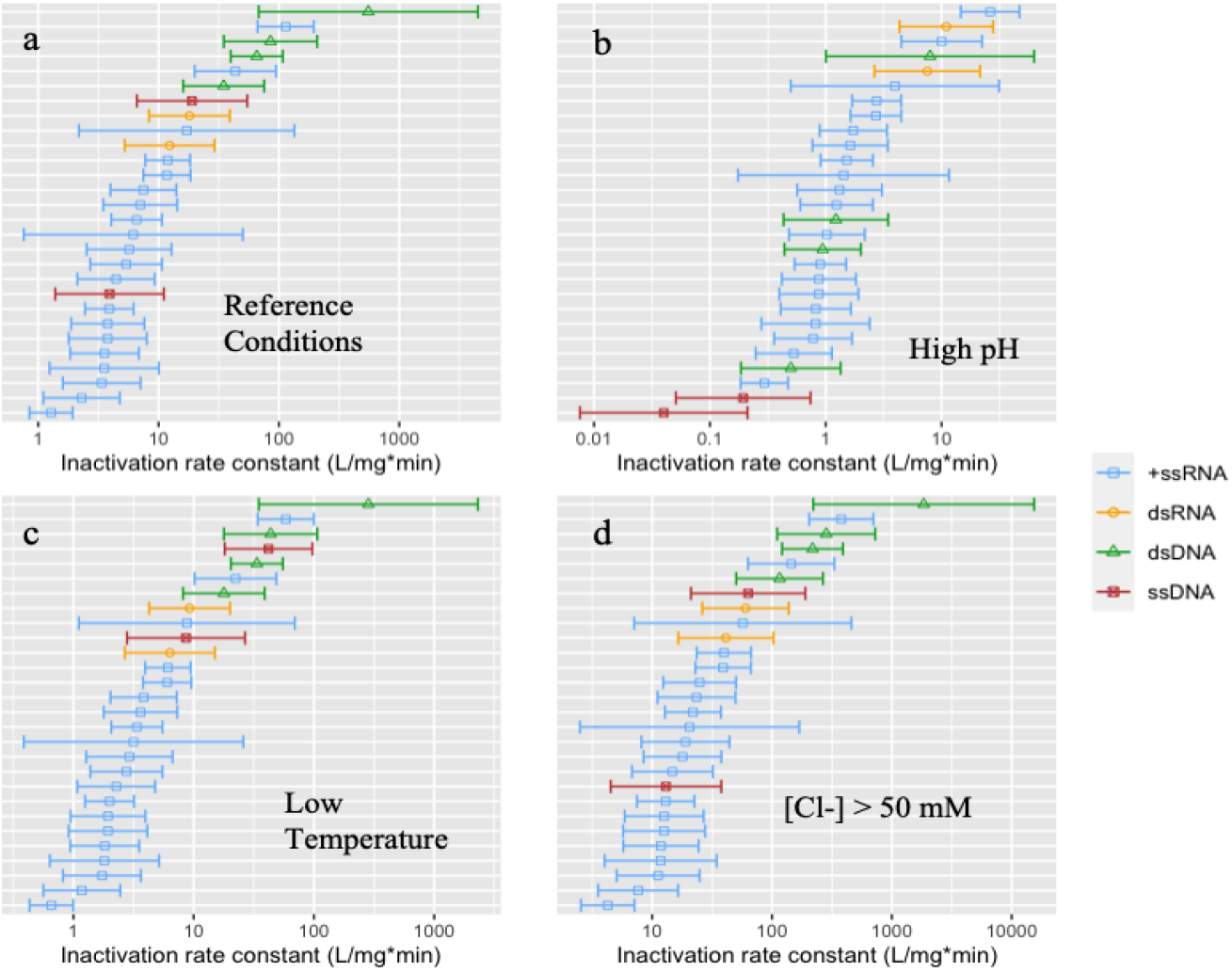
Harmonized virus inactivation rate constants (L/mg*min) and the associate confidence intervals generated by the model under different experimental conditions for viruses with five or more data points in the dataset. Inactivation rate constants are arranged from lowest to highest in each figure. a) Inactivation rate constants at reference conditions (pH = 7.53, T = 20 °C, [Cl-] < 50 mM); b) Inactivation rate constants at elevated pH conditions (pH = 10, T = 20 °C, [Cl-] < 50 mM); c) Inactivation rate constants at low temperature conditions (pH = 7.53, T = 5 °C, [Cl-] < 50 mM); d) Inactivation rate constants under elevated chloride conditions (pH = 7.53, T = 20 °C, [Cl-] > 50 mM).

As noted above, the model includes Baltimore class-specific interaction terms (e.g., dsRNA and pH; ssRNA and temperature), meaning that the predicted rate constants of viruses from different Baltimore classes are impacted differently by changes in experimental conditions. For example, a pH increase from 7.53 to 10 results in a predicted rate constant decreasing 4-fold for (+)ssRNA viruses, 1.6-fold for dsRNA viruses, and 70-fold for dsDNA viruses. This suggests that the relative susceptibility of viruses to chlorine depends on experimental conditions. Model results indicate that echovirus 1 Farouk is more resistant than human adenovirus 5 under reference conditions (pH = 7.53, T = 20 °C, [Cl-] < 50 mM) but human adenovirus 5 is more resistant than echovirus 1 Farouk at high pH (pH = 10, T = 20 °C, [Cl-] < 50 mM). The mechanistic reasons for these virus-specific effects of experimental conditions are likely the result of different virus component reactivities under different solution conditions. For example, the relative reactivities of hypochlorous acid (HOCl) and hypochlorite (OCl-) may differ among virus protein and nucleic acid targets in different viruses. Additionally, the structure of different viruses may change under changing environmental conditions, exposing or hiding reactive functional groups.

Results from the model provide an opportunity to revisit US EPA guidelines for chlorine disinfection of viruses. These guidelines were developed based on hepatitis A virus (HAV) HM175 data from a 1988 study,^19^ and with an incorporated safety factor.^16,50^ At the time of the study, the measured HAV rate constants were relatively conservative compared to data published on other viruses. Our harmonized data, however, suggest that HAV inactivation rate constants are in the mid-range of inactivation rates for human pathogens. We converted the guideline CT values for 4-log_10_ virus reduction to equivalent rate constants to directly compare the guideline data with our model results under environmental conditions relevant to drinking water treatment (see Supplementary Information section S9 for calculation). Although the US-EPA guidelines require the same CT values for pH values between 6 and pH 9 due to HAV results of the 1988 study, our model suggests that pH does affect virus inactivation in this range, and predicts a 3.6-fold increase in CT values for MS2 bacteriophage between pH 6 and pH 9. With respect to temperature, the free chlorine CT requirements represent a sixfold decrease in CT value between temperatures of 0.5 **°**C and 25 **°**C; the statistical model, however, predicts only a threefold decrease in CT value between these temperatures. For every condition described in the US EPA guidelines, our model predicts that at least five of the viruses in our dataset do not achieve 4-log_10_ inactivation under the EPA required CT values (Table 1), with even more viruses not achieving 4-log_10_ inactivation at pH 9 and temperatures above 20 **°**C. The statistical model results suggest that current CT guidelines may not achieve 4-log_10_ inactivation for many viruses under elevated temperatures and pH values. This confirms a concern raised in the most recent EPA Six-Year Review that current regulations may be insufficiently conservative to achieve CVB5 removal at these conditions.^16^

**Table 1.**
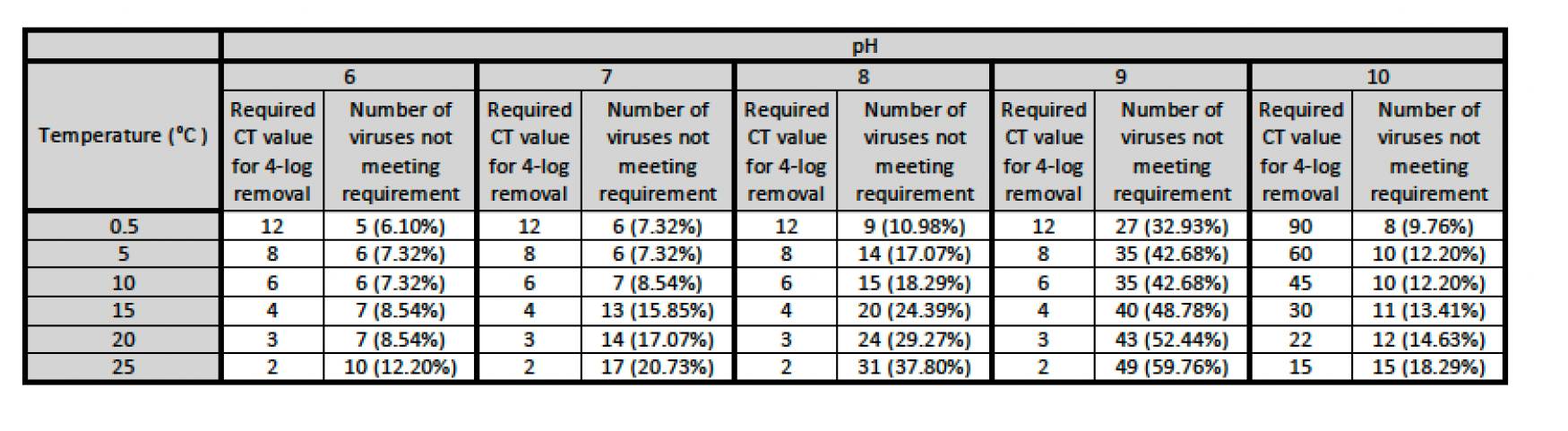
Number and percentage of viruses (out of 82 total viruses) predicted by our model not to meet the requirement for 4-log_10_ removal under EPA required CT values.

#### Correlation of Error within Papers

Random effects (i.e. the correlated within-paper error) played a large role in the predicted chlorine inactivation rate constants, indicating that inactivation rates for different viruses are systematically higher or lower when they are reported in the same publication. The standard deviation of the random effects was 0.448 log_10_ units (Figure S11), meaning that a hypothetical paper drawn from the population has a 32% chance of having a rate constant be biased by at least ±2.8 folds (= 10^0.448^). Note that 32% represents the probability that a random variable drawn from a Normal distribution is more than 1 standard deviation above or below the mean. The adjusted Intraclass Correlation Coefficient (ICC), which refers to the correlation between rate constants calculated from the same paper, is 0.700 (95% CI:.579-.780). Based on this, 70.0% of variance unexplained by the fixed effects is explained by the random effects.

A mean random effect of exactly zero, which is found for 8 papers, indicates that a paper reported inactivation data for a virus that was not studied in another paper. In these cases, the model could not distinguish between virus effects, random effects, and unexplained residual error, and thus the mean random effect of the publication ID was estimated to be zero. A small but non-zero random effect indicates that the paper studied viruses that had been studied elsewhere at commonly studied conditions and produced results largely similar to those produced by other papers. A large random effect indicates that the paper studied viruses that had been studied elsewhere at commonly studied conditions and produced results largely different to those produced by other papers. Papers exhibiting large random effects may be systematically biased, possibly because certain experimental details were not accurately reported (e.g., room temperature, etc.) or were not captured in the model (e.g., reactor stir speed).

#### Environmental Implications

This comprehensive systematic review of chlorine inactivation of viruses represents a unique and important resource for scientists and practitioners. Previous systematic reviews of chlorine inactivation of viruses focused on a narrow range of water conditions^51^ or on virus families commonly emphasized in water treatment, including enteroviruses, adenoviruses, and caliciviruses^30,49,51^. By reviewing chlorine inactivation across all studied viruses and conditions, we were able to build a statistical model that filled in important experimental condition gaps and harmonized data for all viruses under similar environmental conditions. To aid researchers in their use of our dataset, we have produced a website to facilitate the use of our predictive model (https://virus-inactivation.shinyapps.io/chlorine_virus_model/). This website allows for immediate predictions of inactivation rate constants for all studied viruses under a variety of conditions.

Our dataset highlights the lack of any high-quality chlorine data for several viruses important to human health. Whereas the extensive available enteroviruses data highlights a wide range of inactivation rate constants for closely related species and strains, more data is needed for closely related strains of other human viruses (e.g., rotaviruses, caliciviruses) to understand if they too can have highly varied rate constants. Recent outbreaks and pandemics of viruses not closely associated with drinking water have highlighted the need for high-quality disinfection data for a broader range of viruses, including those without known fecal-oral transmission routes and those that infect animals, and more chlorine data and further modeling is necessary to assess removal of these viruses.

The systematic review and model identified several study details that could inform future studies of virus inactivation by chlorine. Specifically, multiple data points from otherwise excellent studies were excluded from the systematic review because the study did not characterize chlorine decay, making it impossible to determine chlorine exposure. To provide external validity and thus measure random effects, future studies of virus reactions with chlorine should include controls with reference water treatment conditions (*i.e.* room temperature, neutral pH, low chloride concentrations) and include a widely studied reference virus. Potential reference viruses include MS2 bacteriophage, CVB5 Faulkner, human adenovirus 2, echovirus 1 Farouk, and echovirus 11 Gregory as these have all been studied in at least five papers and under a variety of conditions.

The statistical model is a valuable resource for selecting surrogates for human waterborne viruses. Process surrogates are commonly used to stand in for all human viruses to test water treatment processes with site-specific reactors and waters. Likewise, experimental surrogates are used by researchers to estimate chlorine inactivation rates of specific human viruses that are difficult to culture (e.g., murine norovirus used in lieu of human norovirus^52^). For chlorine treatment, surrogates should be equally or more resistant to chlorine than human viruses of interest. MS2 bacteriophage is the most common surrogate because it is structurally similar to a number of important human enteric viruses, relatively easy to culture, and safe to work with. Our results suggest that although MS2 bacteriophage has one of the lowest bacteriophage inactivation rate constants at reference conditions, it is not a conservative surrogate for many human viruses with free chlorine. It is an especially poor surrogate for enteroviruses, with 40 out of 43 studied enteroviruses having predicted rate constants lower than MS2 bacteriophage under all environmental conditions. CVB5 Faulkner had the fifth-lowest predicted rate constant of all viruses included in the systematic review, and the lowest predicted rate constant of all viruses that represented at least five data points, suggesting that it is a conservative surrogate for human viruses overall. Whereas CVB5 could serve as a conservative surrogate for human viruses with free chlorine, its BSL2 characterization means it cannot be easily used in all labs. Ideally, a bacteriophage or animal virus would be identified that is at least as conservative as CVB5 and easy for laboratories to work with. We note that experimental conditions should be considered in the selection of conservative surrogates for chlorine disinfection as some viruses are relatively more conservative under certain conditions than others.

There are limitations to the literature review and statistical modeling. We built our model with rate constants calculated using only the first-order portion of inactivation curves, and then assumed that these inactivation rates could be applied to 4-log_10_ inactivation, although some studies have shown significant tailing before 4-log_10_ inactivation. If this assumption is incorrect, viral inactivation would be slower than predicted by our model, and even more viruses than presented in Table 1 would not achieve 4-log_10_ inactivation under required CT values. One limitation of the statistical model is that it cannot draw distinctions below the taxonomic level of virus name, even when such distinctions are known to exist. These limitations can be addressed as more researchers sequence the viruses they study, enabling us to narrowly distinguish viruses. Additionally, some of the assumptions in our model are biased by our limited dataset, including that we assumed viruses in the same Baltimore class would respond similarly to changes in environmental conditions. As more data becomes available, the model can be updated to make finer distinctions. Ultimately, this literature review and statistical analysis provide tools for researchers and practitioners to identify new research directions and evaluate treatment processes.

## Supporting Information

The following files are available free of charge. Files S1-S10 are available in a word document.

S1. Supplementary Methods for Systematic Review and Data Extraction

S2. Details on Calculation of Inactivation Rate Constants

S3. Variable Description and Derivation

S4. Model Selection

S5. Examination of Limitations in the Dataset

S6. Data Comparisons under Narrowed Environmental Conditions

S7. Coefficients for Final Model

S8. Analysis of Relative Inactivation Rates across Baltimore Classes

S9: Calculations for Assessment of EPA standards

S10: Random Effects

S11: Predicted Inactivation Rate Constants of all Viruses under Reference Conditions (csv)

## Acknowledgements

This work was supported by National Science Foundation project #2015187. MC was supported by a National Science Foundation Graduate Research Fellowship. KL was supported by the Jack Kent Cooke Foundation and the Summer Undergraduate Research in Engineering Program.

## For Table of Contents Only

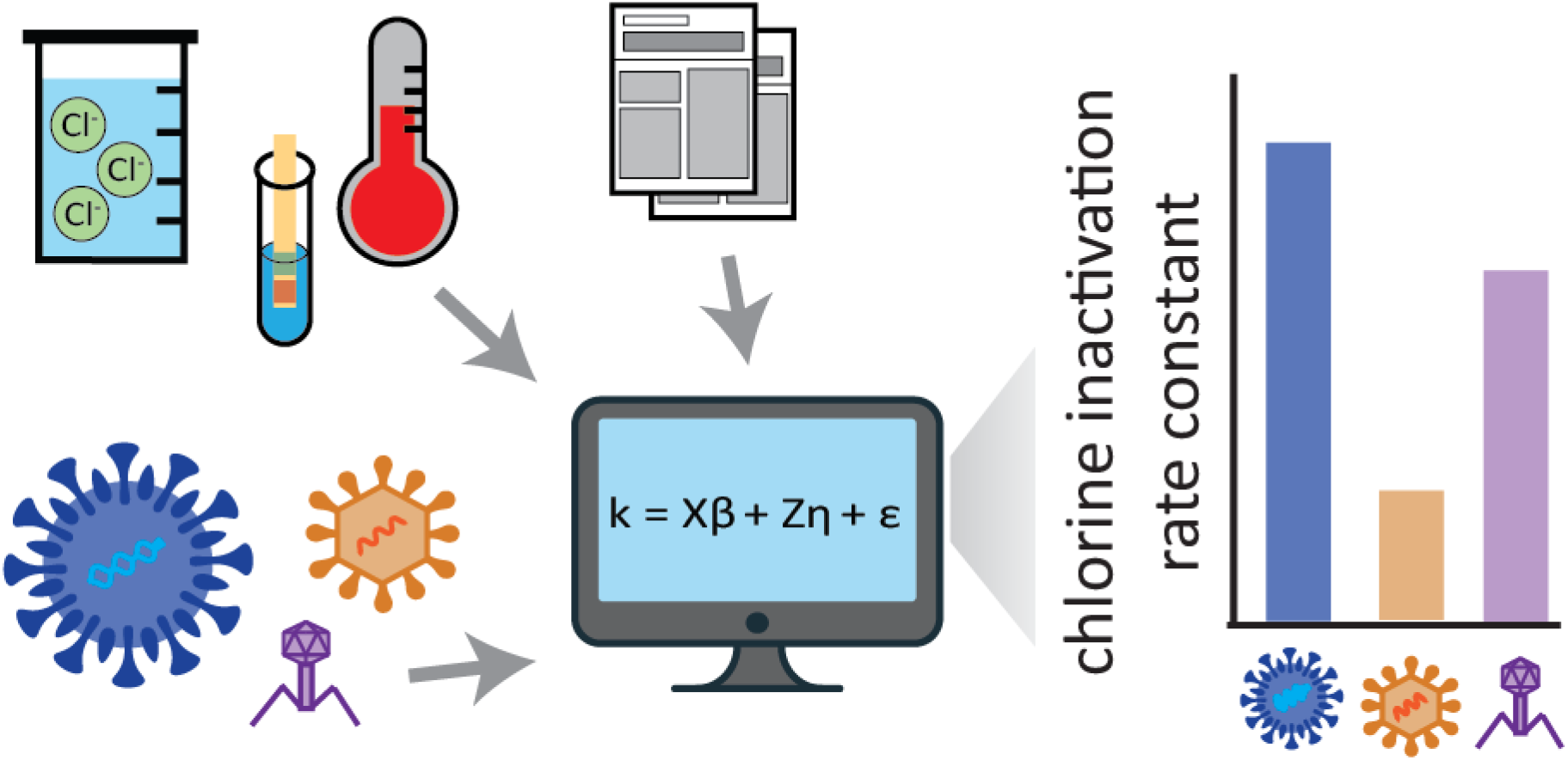

