## Supporting Information 1-10 for "Linear Mixed Model of Virus Disinfection by Chlorine to Harmonize Data Collected Across Broad Environmental Conditions"

### Supplementary Information

#### **S1. Supplementary Methods for Systematic Review and Data Extraction**

**Details on Inactivation Criteria**

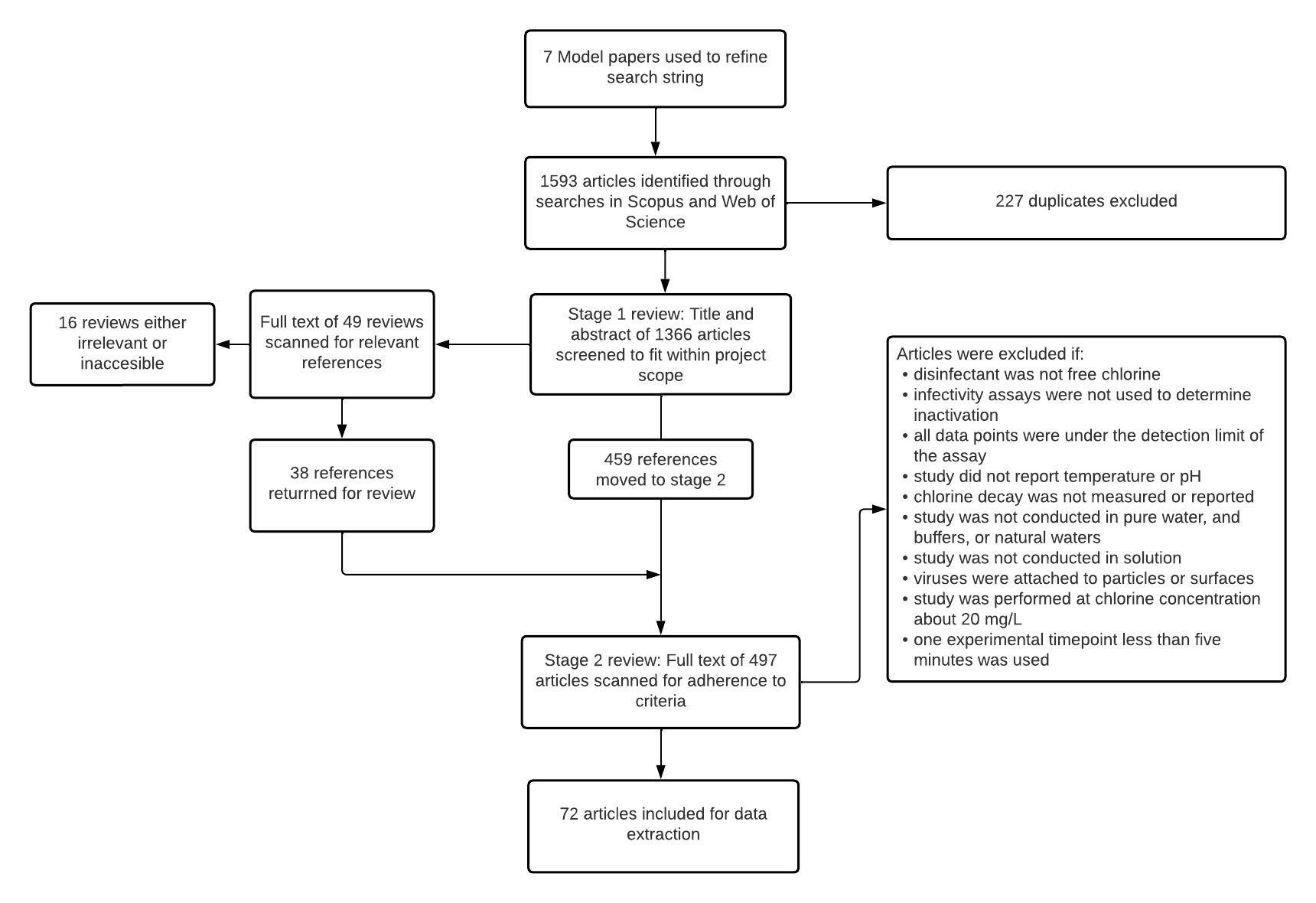

**Figure S1**. Flowchart for systematic review. The two-step review resulted in 72 articles.

Prior to the first stage of the systematic review, we selected seven model publications to develop study inclusion criteria (Figure S1). Both Scopus and Web of Science databases were searched with test search strings until the search results from both databases contained the seven model papers. The resulting search string was applied to both Web of Science and Scopus on January 13, 2021: “((chlorine OR free chlorine OR hypochlorous acid OR HOCl OR hypochlorite OR bleach) AND (inactivat* OR disinfect* OR degradat* OR kinetic* OR infectivity OR decontaminat* OR sanitize) AND (*virus OR virus OR viral OR phage OR bacteriophage))”. Duplicates were flagged by Rayyan software and manually removed.

In the first stage, titles and abstracts were independently scanned by two reviewers to determine 1) if the article was written in English and 2) if the study involved virus inactivation by chlorine in solution. An article was excluded at this stage only if both reviewers agreed that it did not meet these criteria. Review articles identified in stage one were read for relevant references that had not already been identified. In the second stage, the full article text was read by one of three reviewers, and articles were included for data extraction based on specific criteria that supported high-quality chlorine inactivation data. Broadly, we set criteria to ensure that chlorine concentration and virus infectivity were accurately reported and chlorine concentrations were relevant for water and wastewater treatment systems.

Articles were included if:

1. The disinfectant was free chlorine.
2. Culture-based assays were used to determine inactivation, with the exception of assays done using non-microbial organisms, such as plants or abalone. Non-microbial assays were excluded because they had much smaller sample sizes. Because chlorine damages both the genome and the protein of viruses^1^ there is no broadly applied correlation between inactivation rate and genome decay. Loss of PCR signal may occur without inactivation, and inactivation may occur while PCR signal remains intact. Therefore, papers using non-culture based assays were excluded.
3. The study reported temperature and pH. Due to the previously demonstrated impact of temperature and pH on chlorine inactivation of viruses, we concluded that, in a study that did not report these parameters, it would be impossible to contextualize differences between viruses.
4. Chlorine decay was measured or reported. Studies were included if the chlorine concentration was said to be constant or to decay by less than 20%, if the initial and final chlorine concentrations were reported, or if chlorine decay parameters (such as first-order chlorine decay rate) were reported. Additionally, studies were included if reported CT values were calculated using a model such as the Efficiency Factor Hom model, which incorporates chlorine decay^2,3^. Studies were also included if extensive virus purification procedures were undertaken, along with glassware processing to remove chlorine demand. Extensive purification was defined as purification done with a gradient method, including sucrose or cesium chloride, column purification, or HPLC.
5. Study was conducted in pure water, buffers, natural waters, or wastewater that had undergone secondary treatment. Studies conducted in agricultural wash waters were included. Studies conducted in human secretions or in raw sewage were not included. While no turbidity limit was set, when articles included datasets with varying turbidity, only the lowest turbidity values were selected.
6. Study was conducted in solution.
7. Viruses were not attached to particles or surfaces. Studies where viruses were deliberately aggregated were not included.
8. Study was performed at chlorine concentrations below 20 mg/L. This limit is close to the typical chlorine concentration used in drinking and wastewater treatment. At higher concentrations, kinetics may differ.
9. The study included experimental timepoints under five minutes.
10. The study reported first or second order rate constants, CT values for various log_10_ inactivation rates, N/N0 values at specific times, or included a graph where one of these parameters could be extracted. We only required one time point or CT value in addition to a well-characterized initial sample.
11. The study was performed on a virus with a publicly available genome. Viroids and general virus groupings, including “somatic coliphages” were excluded.

#### **S2: Details on Calculation of Inactivation Rate Constants**

We determined the second-order rate constant k from the equation:

$$\ln\left( \frac{N}{N_{0}} \right)=-k(CT)$$

by linear regressions of ln(N/N_0_) and CT. Whenever possible, we calculated CT values directly. from the reported chlorine concentrations and reaction times. When CT values were directly reported or minimal chlorine decay information was reported, three other methods were used (Table S1).

**Table S1.** Methods of calculating CT values from papers

| Case | Method of Reporting Inactivation and Chlorine | Equation Used to calculate CT values | Paper References |
| --- | --- | --- | --- |
| 1 | Timepoints and concentrations provided, chlorine assumed constant | $CT=C*t$ | ^4–37^ |
| 2 | Chlorine decay reported and first-order chlorine decay rate constant provided | $CT=\frac{C_{0}}{k_{d}}(1-e^{-k_{d}t})$ ^38^ | ^39–42^ |
| 3 | Chlorine concentrations reported throughout experiment | ${CT}_{n}=\sum_{0}^{n-1} CT+\frac{C_{n+1}+C_{n}}{2}*(t_{n+1}-t_{n})$ | ^43–51^ |
| 4 | CT values directly provided | CT values used from paper | ^7,8,16,38,43,52–74^ |

*Case 1*

When manuscript text suggested that chlorine concentrations were stable through the experiment, CT values were calculated by multiplying timepoints by the reported chlorine concentration. Chlorine concentrations were considered stable if the manuscript stated that 1) the concentration decreased by less than 20% over the course of the experiment or 2) the described experimental conditions suggested the experimental solutions had a low chlorine demand, namely that virus stocks were purified using gradient purification, column purification, or High-Performance Liquid Chromatography.

*Case 2*

In a number of cases a first order rate constant for chlorine decay was reported. In these cases, CT values were calculated at each experiment time t according to the following equation^38^:

$CT=\frac{C_{0}}{k_{d}}(1-e^{-k_{d}t})$

Where C_0_ is the initial chlorine concentration and k_d_ is the first-order chlorine decay rate.

*Case 3*

In the cases where chlorine concentrations changed through the experiment and were specified, CT values for each dose were determined based on the following relationship:

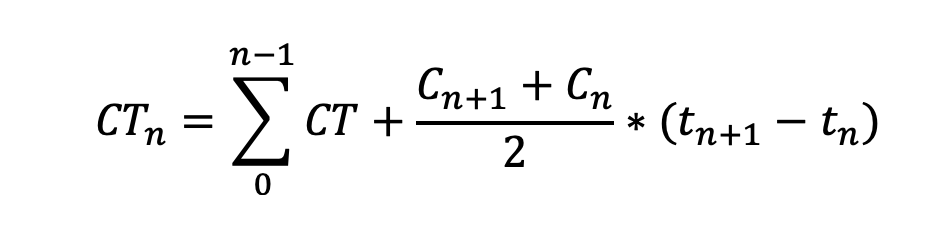

where CT_n_ is the CT value at any time point t_n_.

*Case 4*

In a number of cases insufficient experimental data was provided in figures and tables to calculate the CT values externally. In these cases, we used CT values reported in the manuscripts to directly calculate k values. The CT values reported in the manuscripts had been calculated using the approaches described above as well as a variety of other methods, including the Efficiency Factor Hom (EFH) model. We note that although the EFH model is widely used in cases where both chlorine and viruses are decaying, we elected not to apply it in our study as the model parameters are not directly interpretable and it requires data missing from many of the included studies.

When data was presented differently for experiments reported in the same paper, multiple approaches were used for the same paper. For all cases, when rate constants calculated by the two reviewers differed by greater than ten percent, the calculations were reviewed and corrected if a discrepancy was identified. In all cases, the average of the two rate constants collected by the two reviewers was used for the model.

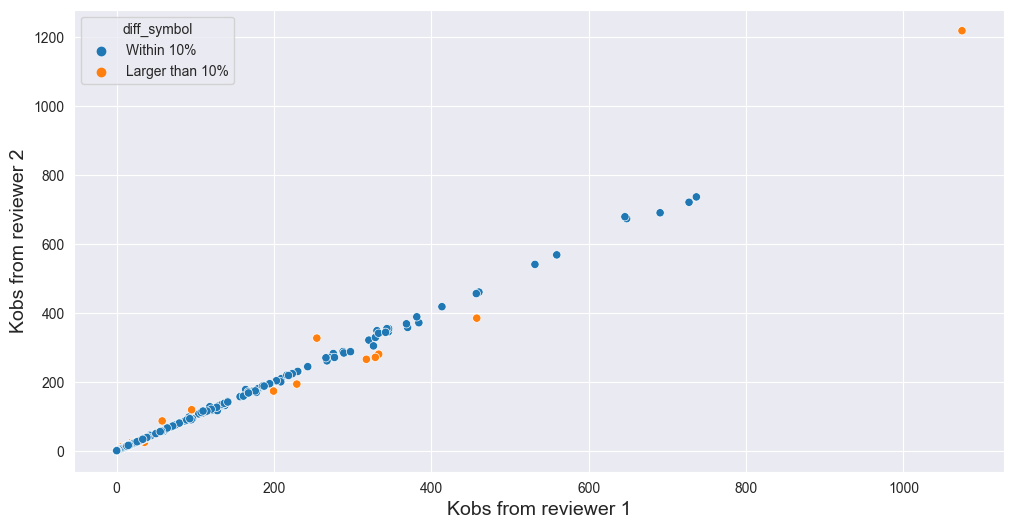

**Figure S2** Lin’s Concordance between final rate constants as calculated by two reviewers. Data points that differed by >10% between reviewers are marked in orange.

#### **S3: Variable Description and Derivation**

*Water type.* We divided water type into four categories: buffer solutions, natural or treated waters, ultrapure water, and high organics solutions. This is because different study solutions have been shown to impact chlorine inactivation. Buffer solutions were either purchased or ultrapure buffers augmented with salts, including acetate, borate, calcium, carbonate, magnesium, phosphate, phosphate carbonate, phosphate saline, phthlate. Low organics natural or treated waters include surface water, tap water, and treated groundwater and surface water. Ultrapure water is distilled, miliQ, or water that has been pH adjusted with only NaOH or HCl. Natural High organics natural or treated water includes wastewater and agricultural waters used to wash lettuce. While differences exist within buffer categories that could impact inactivation, the small number of data points for some categories (e.g. acetate) raised risks of overfitting if they were incorporated into the model.

*Chloride level.* Solution classification as high or low chloride was complicated by the broad use of Phosphate Buffer Saline, often abbreviated as PBS. Solutions with many different chloride concentrations are abbreviated as PBS, some of which (e.g., Dulbecco’s PBS) have greater chloride content than our threshold, and others of which have lower chloride content than our threshold. Solutions simply described as Phosphate Buffer Saline or abbreviated as PBS were assumed to have low chloride if no other information was provided.

*Virus purity.* As virus purification (among other experimental measures) prevents chlorine decay, we therefore defined purification level as an experimental variable. Purification level was divided into three categories: high, medium, and low. Low purification was defined as studies when no purification method was mentioned, or when only low levels of centrifugation were used to eliminate cellular debris. High purification was defined as studies that used gradient purification, column purification, or high performance liquid chromatography. Medium purification was defined as intermediate between these two levels, including studies that used filtering through .22 uM filters, PEG precipitation, chloroform extractions, or ultrafiltration. As virus purification technologies are complex and may occur in different orders in different studies, no attempt was made to include purification in the dataset in more granularity.

*Virus taxonomy.* Virus taxonomic information was extracted from the International Committee on Taxonomy of Viruses (ICTV) and is based on the 2021 taxonomic release^75^. When no information aside from the virus name (e.g., Echovirus 1) was provided, assumptions were made to find the corresponding genetic information. If a virus was described as a laboratory strain, it was assumed to be the strain sold by ATCC. If another paper on the same virus had been published during the same year by the same researcher, those viruses were assumed to be the same strain. When no assumptions were possible, the virus was assumed to be a reference strain listed on the ICTV website. The variable “virus name” was defined as the most specific taxonomic unit at which genomes could be found to differentiate viruses. The genomes of each virus with reported inactivation rates were found from NCBI, or from ATCC if the strains were ATCC strains. The genome length was found by calculating the nucleotide length of each downloaded genome file.

*Virus Morphology.* Virion features were identified for the viruses included in the extracted data set. These features include particle diameter, the presence of an envelope, the presence of a tail, and particle symmetry. As virus diameter varies (polymorphism), when diameter ranges were reported, the average diameter was calculated for each virus. This was also done for non-spherical viruses, including rhabdoviruses. When diameters of reported strains were not available, diameters were assumed to be those of a closely related virus (e.g., GA bacteriophage was assumed to have the same diameter as MS2).

*Other study data.* Each article was reviewed to determine the corresponding author and year of publication. When no corresponding author was designated, the corresponding author was assumed to be the author in whose lab the experiment was conducted, based on author descriptions. When no descriptions were provided, the corresponding author was assigned as the person who had served as corresponding author on the highest number of other papers included in the review.

**Table S2.** Subset of experimental variables tested in the model

| Experimental Variables | Included in the final model? | Literature References/Justification |
| --- | --- | --- |
| Temperature | Yes | ^33,38^ |
| pH | Yes | ^38^ |
| ΔpH^2^ | Yes | HOCl and OCl- have different kinetics with viruses and HOCl relates nonlinearly to pH^19^ |
| High Chloride | Yes | ^24,25,76^ |
| Water Type | No | ^31,37^ |
| α_HOCl_ (Proportion of free chlorine as HOCl) | No | HOCl and OCl- have different kinetics with viruses and HOCl relates nonlinearly to pH^19^ |
| Purification | No | ^77^ |
| High chloride x pH | Yes | ^76^ |
| Publication Variables |  | Standard variables to test in linear mixed model |
| Publication Identifier | Yes |  |
| Corresponding Author Identifier | No |  |
| Year of Publication | No |  |
| Interactions (subset of those tested) |  | Identified using residuals testing (see Figure S4.3) |
| High chloride x pH | Yes |  |
| dsDNA : pH | Yes |  |
| dsRNA : pH | Yes |  |
| ssDNA : pH | Yes |  |
| (-)ssRNA : pH | No |  |
| ssDNA : temp | Yes |  |
| DNA : ΔpH^2^ | Yes |  |

**Table S3.** Virus variables tested in the model

| Virus Variables Tested | Included in the final model? |
| --- | --- |
| Virus Name | Yes |
| Species | No |
| Genus | No |
| Family | No |
| Baltimore Class | No |
| Diameter | No |
| Genome Length | No |
| Tail | No |
| Symmetry | No |
| Envelope | No |

#### **S4: Model Selection**

To find the model with the lowest AIC, we began with a model including only virus name and then tested various experimental features known to impact virus inactivation. A model with virus name as the only virus feature produced an AIC of 1099.94 (R^2^ =.463), and including temperature and pH in the minimal model lowered the AIC to 953.91 (R^2^ = 0.580). We next tested both publication ID and corresponding author as random effects. Including publication ID as a random effect lowered the AIC to 744.61 (R^2^ = 0.804), and the addition of corresponding author as a random effect did not lead to a further AIC reduction. High chloride lowered the AIC to 711.57 (R^2^ = 0.818). The inclusion of the buffer as a fixed effect raised the AIC to 7435.10 (R^2^ =.802). Including the purification level of the virus mixture raised the AIC, as did inclusion of the alpha value. Year of publication similarly raised the AIC. Finally, the interaction between pH and high chloride lowered the AIC to 694.44 (R^2^ = 0.825). Ultimately, the M1 model included virus name, temperature, high chloride, and pH as independent variables, as well as the interaction between high chloride and pH, and publication ID as a random effect.

We also incorporated the different responses of viruses to experimental conditions into model selection through the interaction of Baltimore classes and temperature, pH, and ΔpH^2^. A residuals analysis of M1 broken down by Baltimore class, temperature, and pH identified additional trends in the data (Figure S6). Based on these trends we tested ΔpH^2^ as an independent variable, along with added interactions between individual Baltimore classes and temperature, pH, and ΔpH^2^. For each of these variables, we compared multiple options to find the best way to incorporate them into the model. Starting with M1, we sequentially tested the interaction of Baltimore class and pH. We tested Baltimore class in the order of how many data points in the study were from each Baltimore class (dsDNA > dsRNA > ssDNA > (-)ssRNA). We did not test (+)ssRNA interactions, as this was the Baltimore class of the reference virus MS2 bacteriophage; therefore, any Baltimore classes not found to have a significant interaction with Baltimore class were grouped by the model with (+)ssRNA. We included each Baltimore class tested in the model if it lowered the AIC. For pH, the interactions of dsDNA, dsRNA, and ssDNA with temperature lowered the AIC to 622.95 (R^2^ = 0.851). To check if the model improvement was attributable to all Baltimore classes, we also tested M1 non-sequentially with the interaction between all Baltimore classes and pH and found that the AIC dropped to 624.93 (R^2^ = 0.851). Finally, to check if the model improvement was due primarily to different pH effects on DNA and RNA genomes, we tested M1 with the interaction between DNA and pH and found that the AIC dropped to 628.59 (R^2^ = 0.847). We then compared these three models and selected the model including the individual interactions between dsDNA, dsRNA, ssDNA, and pH, because it had the lowest AIC when compared to the non-sequential and DNA interaction models. This model is identified as M2.

To test the interactions between temperature and Baltimore class we started with M2 and followed the same procedure of comparing models produced by the sequentially tested Baltimore class/temperature interactions, the non-sequentially tested Baltimore class and temperature interactions, and the interaction between DNA and temperature. The interaction between Baltimore class and temperature produced a rank deficient model, indicating that not all Baltimore classes were studied at more than one temperature. We found that the only Baltimore class/temperature interaction that improved the AIC was ssDNA and temperature, which lowered the AIC to 607.59 (R^2^ = 0.858). The combination of M2 and the DNA/temperature interaction lowered the AIC to 618.40 (R^2^ = 0.851). Between these two options, the combination of M2 with the ssDNA and temperature interaction was selected and labeled M3.

To test the effect of ΔpH^2^ on the model, we first tested M3 with ΔpH^2^ added and found that it lowered the AIC to 694.35 (R^2^ = 0.860). As with pH and temperature, we then compared models produced by the sequentially tested Baltimore class/ΔpH^2^ interactions, the non-sequentially tested Baltimore class/ΔpH^2^ interactions, and the interaction between DNA and ΔpH^2^. Of these three options, the DNA/ΔpH^2^ interactions produced the greatest model improvement, lowering the AIC to 582.46 (R^2^ = 0.866). The selected model was labelled M4 and includes all terms in M1 as well as ΔpH^2^ and five additional interaction terms (dsDNA and pH, dsRNA and pH, ssDNA and pH, ssDNA and temperature, and DNA and ΔpH^2^).

Due to confounding variables, a linear mixed model of this dataset can include only one virus feature, namely a taxonomy-related or structure-related independent variable. Once the experimental features were selected for M4, we tested the effect of different virus taxonomic units in lieu of virus name on the AIC. Specifically, we built an updated model equivalent to M2 but without virus name (AIC = 902.20, R^2^ = .729) and then tested the impact of adding one of several taxonomic and structural features as independent variables (Figure S5). All of the tested independent taxonomic and structural variables lowered the AIC except for lipid envelope, and the features related to virus taxonomy had greater AIC effects than structural features.

Variables relating to virus structure that lowered the AIC were the presence of a tail (AIC = 898.00, R^2^ = .736) and the capsid symmetry (AIC = 900.42, R^2^ = .731). The presence of an envelope raised the AIC slightly (AIC = 903.39, R^2^ = .732). All of these features were unevenly distributed throughout the dataset; for example, out of 562 data points, only 5 were for viruses that had tails, and only 9 were for viruses that had envelopes. The explanatory power of these variables is therefore limited. Features related to virus size lowered the AIC further, including virus diameter (AIC = 834.49 , R^2^ = .748) and genome length (AIC = 857.85, R^2^ = .732). The inclusion of Baltimore class resulted in an improved AIC (AIC = 821.14, R^2^ = .722). ICTV Taxonomic features further improved the AIC, including species (AIC = 610.59, R^2^ = .829), genus (AIC = 647.27, R^2^ = .803), and family (AIC = 655.56, R^2^ = .792), with narrower taxonomic levels explaining more variance. All other structural and taxonomic variables produced a higher AIC than virus name, despite the large number of columns that virus name added for the data. Ultimately, the virus name was determined to be the best virus feature to include in the model going forward and results from this model are reported throughout the rest of the paper.

The fact that most virus features (tail, capsid structure, diameter, genome length, Baltimore class, family, genus, and species) lowered the AIC and increased the R^2^ value highlights their relevance in virus inactivation by free chlorine.

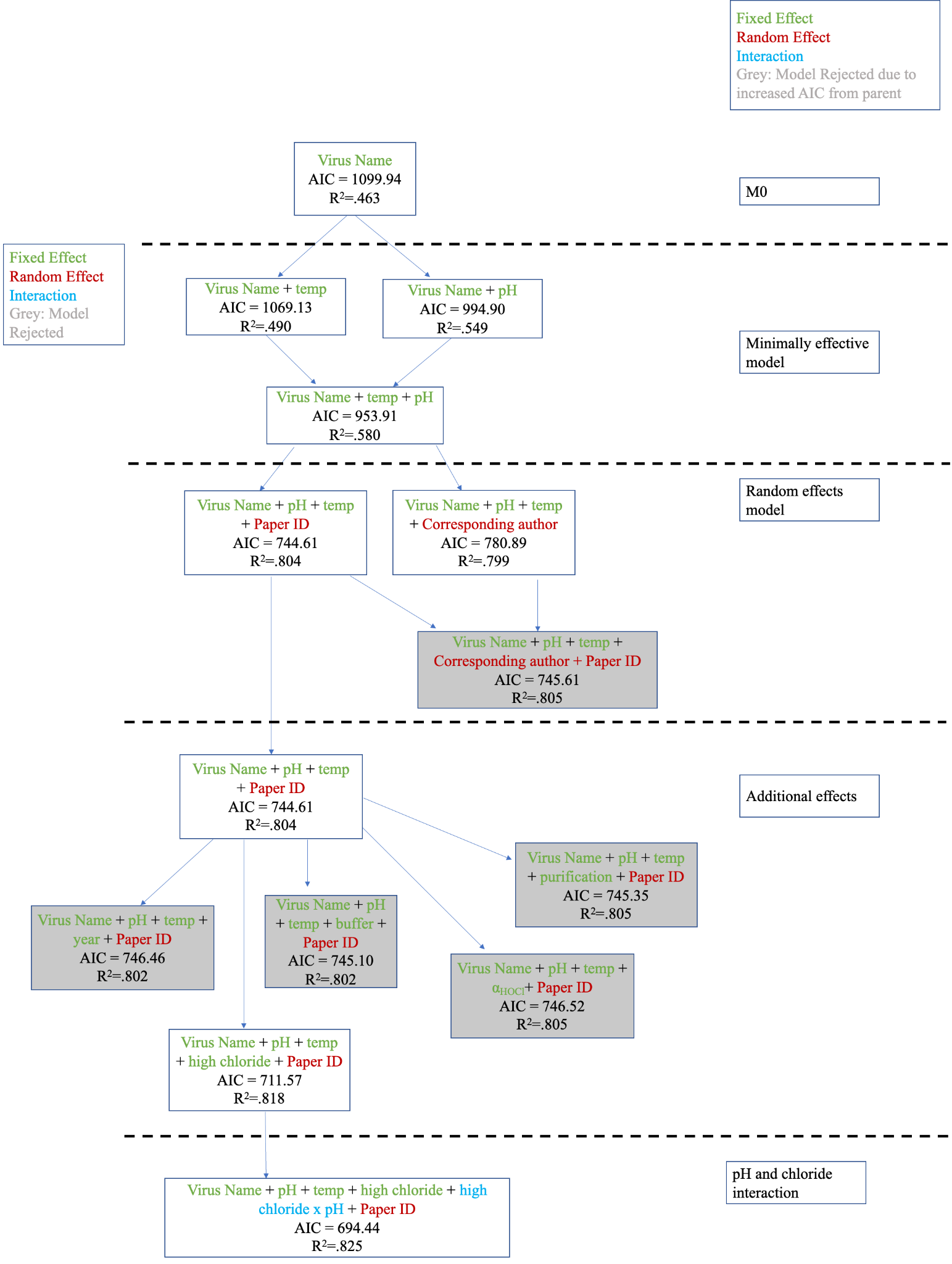

**Figure S3.** Initial model selection figure

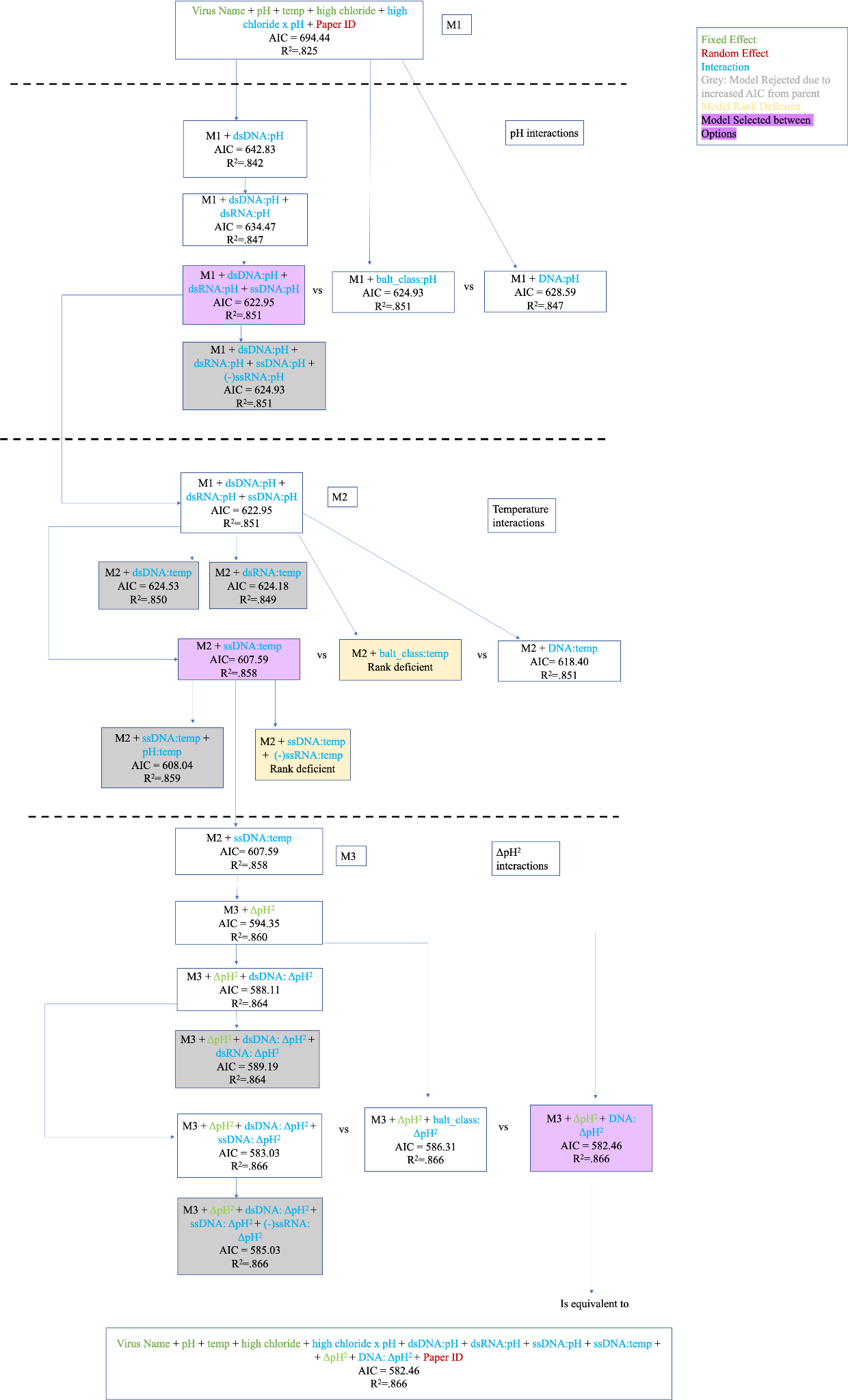

**Figure S4.** Additional interactions model selection figure

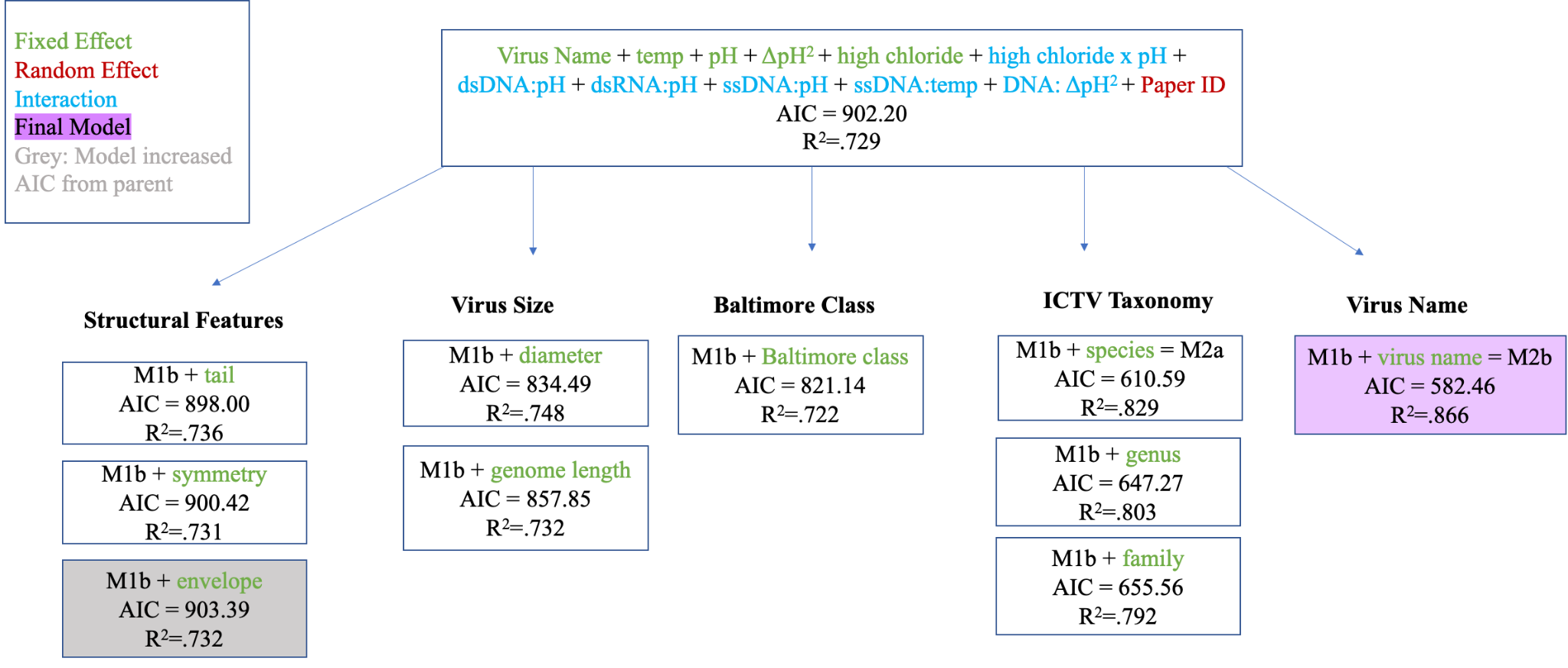

**Figure S5.** Virus taxonomy model selection figure

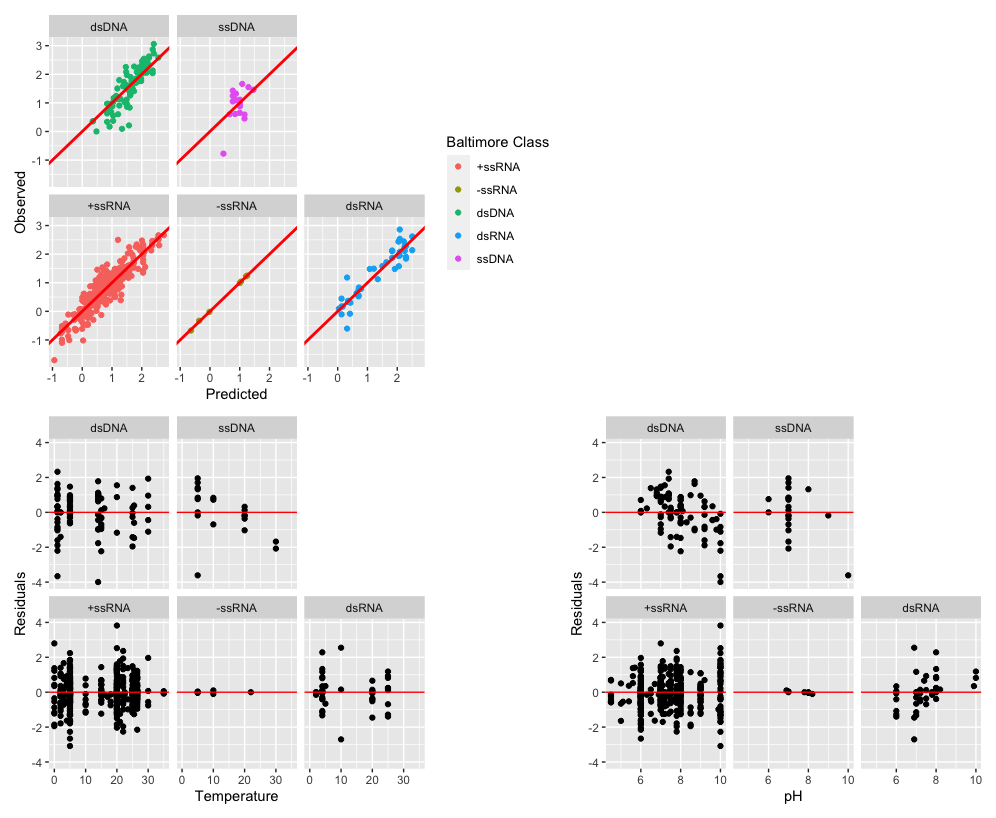

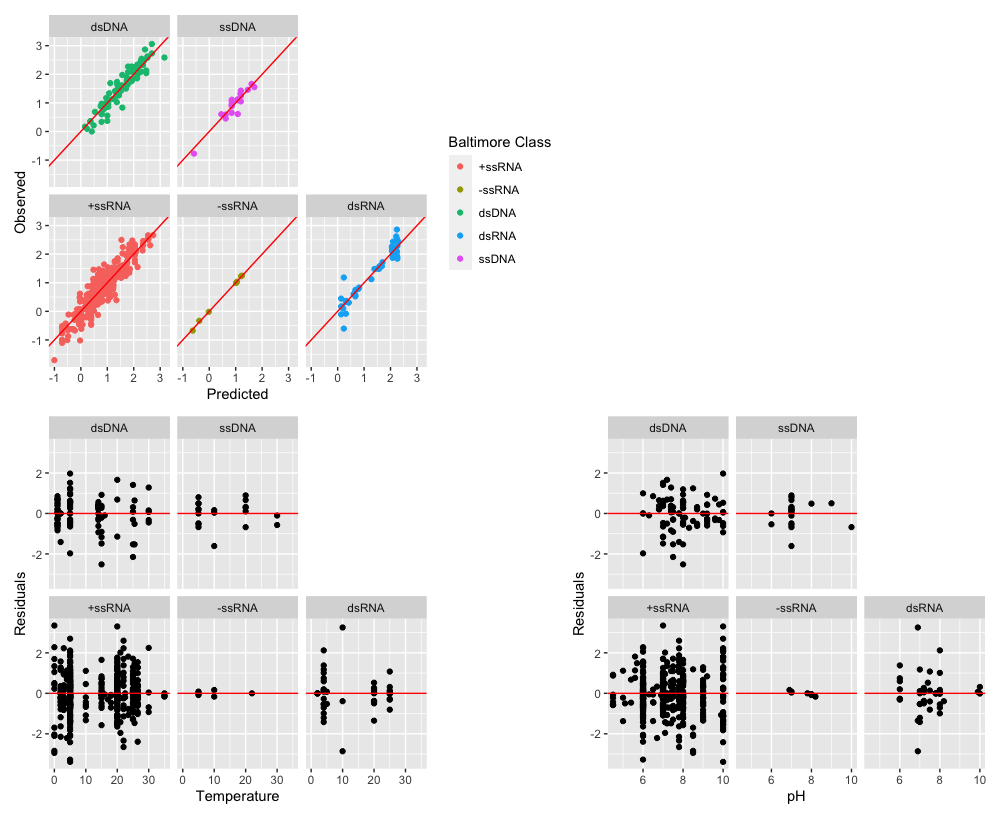

**Figure S6.** Residual analysis comparisons

a) residuals before additional interactions

b) residuals after additional interactions

#### **S5: Examination of Limitations in the Dataset**

**Table S4.** Number of rate constants identified in the review for waterborne viruses of concern, as defined by the WHO

| Viruses | Number of rate constants |
| --- | --- |
| Adenovirus | 63 |
| Astrovirus | 0 |
| Norovirus | 15 |
| Sapovirus | 0 |
| Enterovirus | 297 |
| Hepatitis A virus | 14 |
| Parechovirus | 0 |
| Hepatitis E virus (Orthohepevirus A) | 2 |
| Rotavirus | 32 |
| Orthoreovirus | 6 |

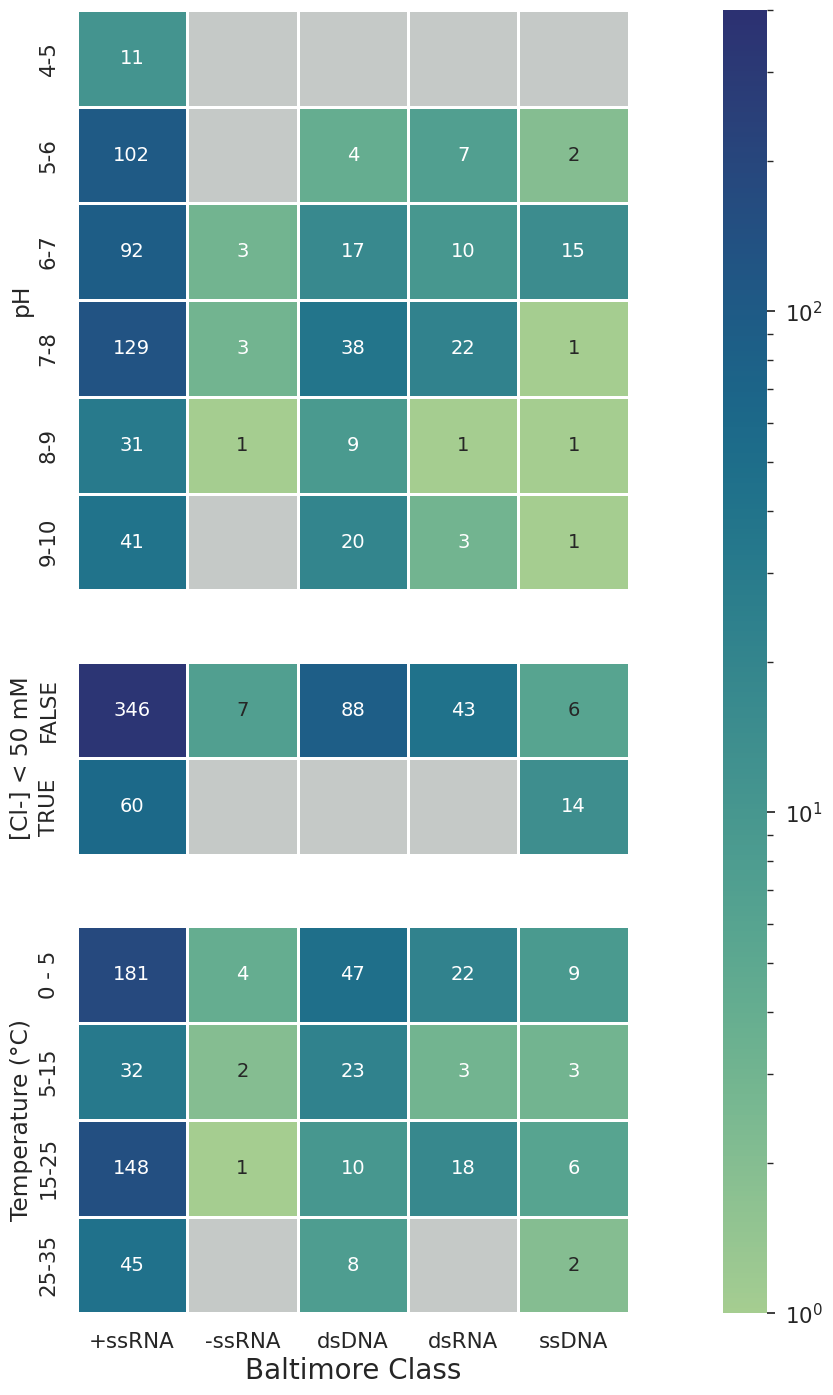

**Figure S7.** Heatmaps showing the number of available data points for viruses in each Baltimore class under a variety of temperature, pH, and chloride levels.

#### **S6: Data Comparison under Narrowed Environmental Conditions**

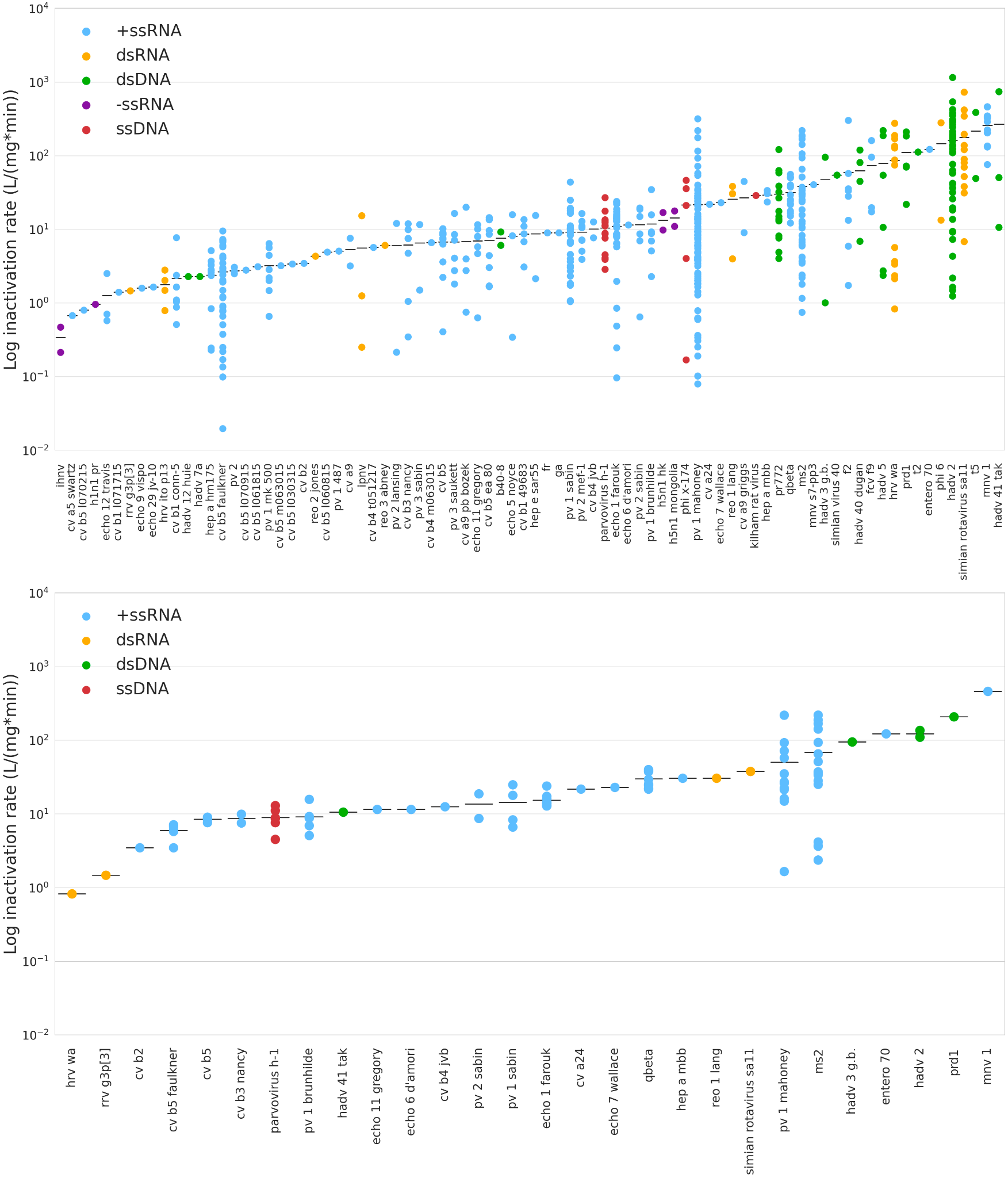

**Figure S8.** a) All inactivation rate constants calculated from the rapid systematic review, ordered from lowest mean to highest mean. b) All inactivation rate constants found at temperatures between 15 and 25 C and pH values between 6 and 7.

#### **S7: Coefficients for Final Model**

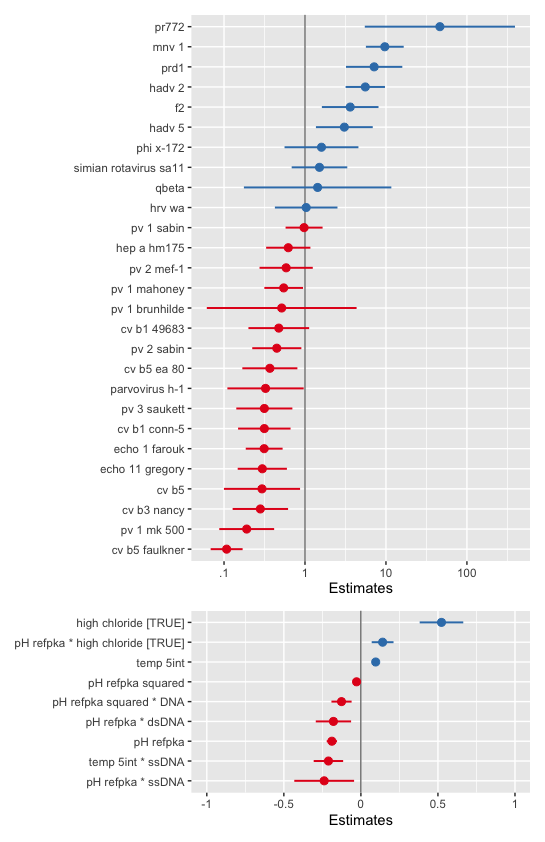

**Figure S9.** a) Coefficients for viruses with five or more data points in the dataset. b) Coefficients for all experimental conditions.

**Table S5.** All Coefficients for Final Model^78^

|  | **Dependent variable** | | |
| --- | --- | --- | --- |
| *Predictors* | *Estimates* | *CI* | *p* |
| (Intercept) | 1.0760 | 0.8893 – 1.2628 | **<0.001** |
| virus_name_strainb40-8 | 0.3616 | -0.1381 – 0.8613 | 0.156 |
| virus name strain [cv  a24] | -0.1979 | -0.8447 – 0.4490 | 0.548 |
| virus name strain [cv a5  swartz] | -0.6727 | -1.2971 – -0.0483 | **0.035** |
| virus name strain [cv a9] | -0.4578 | -0.9507 – 0.0351 | 0.069 |
| virus name strain [cv a9  griggs] | 0.4124 | -0.0676 – 0.8923 | 0.092 |
| virus name strain [cv a9  pb bozek] | -0.2388 | -0.6168 – 0.1392 | 0.215 |
| virus name strain [cv b1  49683] | -0.3204 | -0.6957 – 0.0549 | 0.094 |
| virus_name_straincv b1 conn-5 | -0.4983 | -0.8226 – -0.1740 | **0.003** |
| virus name strain [cv b1  l071715] | -1.0945 | -1.7646 – -0.4244 | **0.001** |
| virus name strain [cv b2] | 0.1505 | -0.6776 – 0.9785 | 0.721 |
| virus name strain [cv b3  nancy] | -0.5481 | -0.8908 – -0.2054 | **0.002** |
| virus name strain [cv b4  jvb] | -0.4196 | -0.8932 – 0.0540 | 0.082 |
| virus name strain [cv b4  m063015] | -0.4211 | -1.0912 – 0.2490 | 0.218 |
| virus name strain [cv b4  t051217] | -0.4847 | -1.1548 – 0.1854 | 0.156 |
| virus name strain [cv b5] | -0.5274 | -0.9979 – -0.0569 | **0.028** |
| virus name strain [cv b5  ea 80] | -0.4294 | -0.7697 – -0.0891 | **0.014** |
| virus name strain [cv b5  faulkner] | -0.9688 | -1.1664 – -0.7712 | **<0.001** |
| virus name strain [cv b5  l030315] | -0.7091 | -1.3792 – -0.0390 | **0.038** |
| virus name strain [cv b5  l060815] | -0.5504 | -1.2205 – 0.1197 | 0.107 |
| virus name strain [cv b5  l061815] | -0.7493 | -1.4193 – -0.0792 | **0.028** |
| virus name strain [cv b5  l070215] | -1.3375 | -2.0076 – -0.6674 | **<0.001** |
| virus name strain [cv b5  l070915] | -0.7935 | -1.4635 – -0.1234 | **0.020** |
| virus name strain [cv b5  m063015] | -0.7355 | -1.4056 – -0.0654 | **0.032** |
| virus name strain [echo 1  farouk] | -0.4851 | -0.7097 – -0.2604 | **<0.001** |
| virus name strain [echo  11 gregory] | -0.5250 | -0.8291 – -0.2209 | **0.001** |
| virus name strain [echo  12 travis] | -0.5998 | -1.0083 – -0.1914 | **0.004** |
| virus_name_strainecho 29 jv-10 | -0.2856 | -0.9100 – 0.3387 | 0.369 |
| virus name strain [echo 5  noyce] | -0.3930 | -0.8093 – 0.0234 | 0.064 |
| virus name strain [echo 6  d'amori] | -0.4766 | -1.1234 – 0.1703 | 0.148 |
| virus name strain [echo 7  wallace] | -0.1756 | -0.8224 – 0.4713 | 0.594 |
| virus name strain [echo 9  vispo] | -0.2994 | -0.9238 – 0.3250 | 0.346 |
| virus name strain [entero  70] | 0.5493 | -0.0975 – 1.1962 | 0.096 |
| virus name strain [f2] | 0.5617 | 0.2116 – 0.9118 | **0.002** |
| virus name strain [fcv  f9] | 0.8420 | 0.2447 – 1.4393 | **0.006** |
| virus name strain [fr] | -0.0202 | -0.7741 – 0.7337 | 0.958 |
| virus name strain [ga] | -0.0189 | -0.7728 – 0.7350 | 0.961 |
| virus name strain [h1n1  pr] | -1.0787 | -2.1484 – -0.0090 | **0.048** |
| virus name strain [h5n1  hk] | 0.3235 | -0.6654 – 1.3125 | 0.521 |
| virus name strain [h5n1  mongolia] | 0.3588 | -0.6302 – 1.3477 | 0.476 |
| virus name strain [hadv  12 huie] | -0.0839 | -0.7088 – 0.5409 | 0.792 |
| virus name strain [hadv  2] | 0.7401 | 0.4970 – 0.9831 | **<0.001** |
| virus name strain [hadv 3  g.b.] | -0.0313 | -0.4996 – 0.4371 | 0.896 |
| virus name strain [hadv  40 dugan] | 0.8265 | 0.2225 – 1.4304 | **0.007** |
| virus name strain [hadv  41 tak] | 0.9106 | 0.4429 – 1.3784 | **<0.001** |
| virus name strain [hadv  5] | 0.4649 | 0.1136 – 0.8161 | **0.010** |
| virus name strain [hadv  7a] | -0.0839 | -0.7088 – 0.5409 | 0.792 |
| virus name strain [hep a  hm175] | -0.2022 | -0.4757 – 0.0713 | 0.147 |
| virus name strain [hep a  mbb] | 0.1023 | -0.3115 – 0.5161 | 0.627 |
| virus name strain [hep e  sar55] | 0.6054 | 0.0433 – 1.1675 | **0.035** |
| virus name strain [hrv  ito p13] | -0.3697 | -0.7864 – 0.0470 | 0.082 |
| virus name strain [hrv  wa] | 0.0166 | -0.3706 – 0.4037 | 0.933 |
| virus name strain [ihnv] | -1.3682 | -2.3564 – -0.3800 | **0.007** |
| virus name strain [ipnv] | -0.6490 | -1.6089 – 0.3110 | 0.185 |
| virus name strain [kilham  rat virus] | -0.5650 | -1.3466 – 0.2167 | 0.156 |
| virus name strain [mnv 1] | 0.9803 | 0.7469 – 1.2137 | **<0.001** |
| virus name strain [mnv  s7-pp3] | 0.5521 | -0.2253 – 1.3294 | 0.164 |
| virus_name_strainparvovirus h-1 | -0.4831 | -0.9548 – -0.0114 | **0.045** |
| virus name strain [phi 6] | 0.8518 | 0.0839 – 1.6196 | **0.030** |
| virus_name_strainphi x-174 | 0.2023 | -0.2543 – 0.6589 | 0.384 |
| virus name strain [pr772] | 1.6672 | 0.7367 – 2.5978 | **<0.001** |
| virus name strain [prd1] | 0.8541 | 0.5060 – 1.2021 | **<0.001** |
| virus name strain [pv 1  487] | -0.7115 | -1.3495 – -0.0735 | **0.029** |
| virus name strain [pv 1  brunhilde] | -0.2858 | -1.2140 – 0.6423 | 0.545 |
| virus name strain [pv 1  mahoney] | -0.2582 | -0.4973 – -0.0190 | **0.034** |
| virus name strain [pv 1  mk 500] | -0.7156 | -1.0548 – -0.3764 | **<0.001** |
| virus name strain [pv 1  sabin] | -0.0063 | -0.2350 – 0.2223 | 0.957 |
| virus name strain [pv 2] | -0.1938 | -1.1546 – 0.7670 | 0.692 |
| virus name strain [pv 2  lansing] | -0.6845 | -1.1644 – -0.2045 | **0.005** |
| virus_name_strainpv 2 mef-1 | -0.2283 | -0.5573 – 0.1008 | 0.173 |
| virus name strain [pv 2  sabin] | -0.3456 | -0.6492 – -0.0421 | **0.026** |
| virus name strain [pv 3  sabin] | -0.2741 | -0.7391 – 0.1909 | 0.247 |
| virus name strain [pv 3  saukett] | -0.4983 | -0.8449 – -0.1518 | **0.005** |
| virus name strain [qbeta] | 0.1580 | -0.7556 – 1.0717 | 0.734 |
| virus name strain [reo 1  lang] | 0.0717 | -0.2892 – 0.4325 | 0.697 |
| virus name strain [reo 2  jones] | 0.0900 | -0.5352 – 0.7151 | 0.777 |
| virus name strain [reo 3  abney] | 0.2376 | -0.3875 – 0.8628 | 0.455 |
| virus name strain [rrv  g3p[3]] | -0.9094 | -1.9798 – 0.1611 | 0.096 |
| virus name strain [simian  rotavirus sa11] | 0.1806 | -0.1633 – 0.5244 | 0.303 |
| virus name strain [simian  virus 40] | 0.4398 | -0.2524 – 1.1320 | 0.212 |
| virus name strain [t2] | 0.4952 | -0.2098 – 1.2002 | 0.168 |
| virus name strain [t5] | 1.6032 | 1.0552 – 2.1512 | **<0.001** |
| temp 5int | 0.0967 | 0.0749 – 0.1186 | **<0.001** |
| pH refpka | -0.1877 | -0.2212 – -0.1542 | **<0.001** |
| high chloride [TRUE] | 0.5227 | 0.3811 – 0.6642 | **<0.001** |
| pH refpka squared | -0.0284 | -0.0512 – -0.0056 | **0.015** |
| pH refpka × high chloride  [TRUE] | 0.1407 | 0.0698 – 0.2116 | **<0.001** |
| pH refpka × dsDNA | -0.1788 | -0.2935 – -0.0642 | **0.002** |
| pH refpka × dsRNA | 0.1704 | 0.0610 – 0.2797 | **0.002** |
| pH refpka × ssDNA | -0.2389 | -0.4330 – -0.0447 | **0.016** |
| temp 5int × ssDNA | -0.2114 | -0.3070 – -0.1157 | **<0.001** |
| pH refpka squared × DNA | -0.1253 | -0.1907 – -0.0598 | **<0.001** |
| **Random Effects** | | | |
| σ^2^ | 0.086 | | |
| τ_00_ _paper_ID_ | 0.201 | | |
| ICC | 0.700 | | |
| N _paper_ID_ | 72 | | |
| Observations | 564 | | |
| Marginal R^2^ / Conditional R^2^ | 0.553 / 0.866 | | |

#### **S8: Genome Length and Diameter Figures**

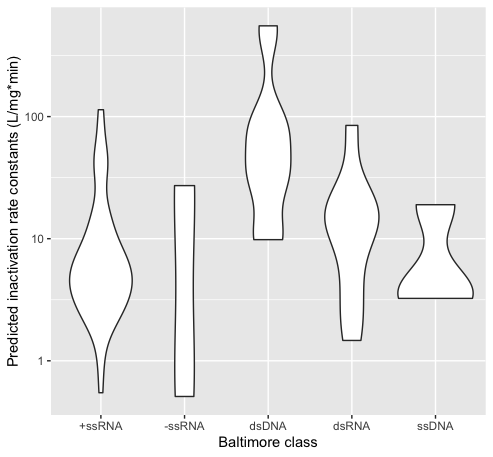

**Figure S10.** Distribution of predicted inactivation rate constants across Baltimore classes

**Table S6.** Diameters and genome length for viruses from each Baltimore class included in the dataset

| Baltimore Class | Diameter (nm) | | | Genome Length (kbp) | | |
| --- | --- | --- | --- | --- | --- | --- |
|  | Mean | Min | Max | Mean | Min | Max |
| (+)ssRNA | 29.8 | 25.8 | 41.6 | 7.03 | 3.47 | 7.69 |
| (-)ssRNA | 105 | 100 | 120 | 13.0 | 11.1 | 13.6 |
| dsDNA | 81.6 | 45 | 131 | 46.9 | 5.24 | 164 |
| ssDNA | 24.7 | 24 | 26 | 5.16 | 4.93 | 5.39 |
| dsRNA | 77.8 | 65 | 80 | 14.4 | 1.42 | 23.6 |

#### **S9: Calculations for Assessment of EPA standards**

**Calculation of required CT value from k value predicted by model**

The equation

$$\ln\left( \frac{N}{N_{0}} \right)=-k(CT)$$

Presented in S2 is rearranged to find the equation

$$CT=\frac{-k}{\ln\left( \frac{1}{10000} \right)}$$

Which provides the CT value to achieve four-log_10_ removal of a virus, given an inactivation rate constant.

#### **S10: Random Effects**

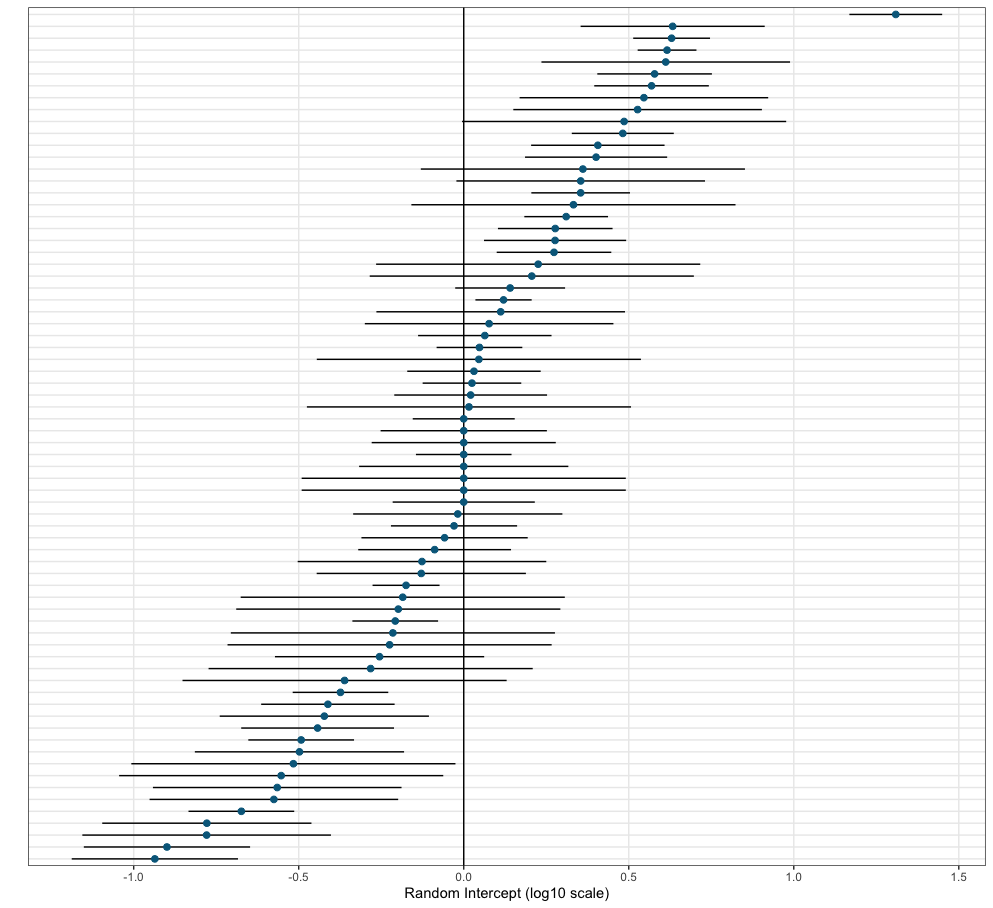

**Figure S11.** Mean of random effects of each paper from the model. Each row represents the random effect from a paper.
